# Gut microbiome dysbiosis during COVID-19 is associated with increased risk for bacteremia and microbial translocation

**DOI:** 10.1101/2021.07.15.452246

**Authors:** Mericien Venzon, Lucie Bernard-Raichon, Jon Klein, Jordan E. Axelrad, Chenzhen Zhang, Grant A. Hussey, Alexis P. Sullivan, Arnau Casanovas-Massana, Maria G. Noval, Ana M. Valero-Jimenez, Juan Gago, Gregory Putzel, Alejandro Pironti, Evan Wilder, Yale IMPACT Research Team, Lorna E. Thorpe, Dan R. Littman, Meike Dittmann, Kenneth A. Stapleford, Bo Shopsin, Victor J. Torres, Albert I. Ko, Akiko Iwasaki, Ken Cadwell, Jonas Schluter

## Abstract

The microbial populations in the gut microbiome have recently been associated with COVID-19 disease severity. However, a causal impact of the gut microbiome on COVID-19 patient health has not been established. Here we provide evidence that gut microbiome dysbiosis is associated with translocation of bacteria into the blood during COVID-19, causing life-threatening secondary infections. Antibiotics and other treatments during COVID-19 can potentially confound microbiome associations. We therefore first demonstrate in a mouse model that SARS-CoV-2 infection can induce gut microbiome dysbiosis, which correlated with alterations to Paneth cells and goblet cells, and markers of barrier permeability. Comparison with stool samples collected from 96 COVID-19 patients at two different clinical sites also revealed substantial gut microbiome dysbiosis, paralleling our observations in the animal model. Specifically, we observed blooms of opportunistic pathogenic bacterial genera known to include antimicrobial-resistant species in hospitalized COVID-19 patients. Analysis of blood culture results testing for secondary microbial bloodstream infections with paired microbiome data obtained from these patients indicates that bacteria may translocate from the gut into the systemic circulation of COVID-19 patients. These results are consistent with a direct role for gut microbiome dysbiosis in enabling dangerous secondary infections during COVID-19.

## Main text

A better understanding of factors contributing to the pathology of coronavirus disease 2019 (COVID-19) is an urgent global priority. Previous reports have demonstrated that severe COVID-19 is frequently associated with specific inflammatory immune phenotypes, lymphopenia, and a generally disproportionate immune response leading to systemic organ failure^1, 2^. Even in mild cases, gastrointestinal symptoms are reported frequently, and recent studies reported that COVID-19 patients lose commensal taxa of the gut microbiome during hospitalization^3–5^. Differences in gut bacterial populations relative to healthy controls were observed in all COVID-19 patients, but most strongly in patients who were treated with antibiotics during their hospitalization^4^. Most recently, COVID-19 patients treated with broad spectrum antibiotics at admission were shown to have increased susceptibility to multi-drug resistant infections and nearly double the mortality rate from septic shock^6, 7^. Furthermore, although initially estimated to be low (6.5%)^8^, more recent studies have detected bacterial secondary infections in as much as 12-14% of COVID-19 patients^9, 10^. However, the causal direction of the relationship between disease symptoms and gut bacterial populations is not yet clear.

Complex gut microbiota ecosystems can prevent the invasion of potentially pathogenic bacteria^11, 12^. Conversely, when the gut microbiota incurs damage, such as through antibiotics treatment, competitive exclusion of pathogens is weakened^13–15^ and potentially dangerous blooms of antibiotic resistant bacterial strains can occur^16, 17^. In immunocompromised cancer patients, blooms of Enterococcaceae and Gram-negative proteobacteria can lead to gut dominations by few or single species^18–21^. Such gut domination events are dangerous to these patients because they are associated with increased risk of translocation of antibiotic resistant bacteria from the gut into the blood stream^18^. Bacterial co-infection can also cause life-threatening complications in patients with severe viral infections^7, 8, 22^; therefore, antibacterial agents were administered empirically to nearly all critically ill suspected COVID-19 patients since the incidence of bacterial superinfection was unknown early during the pandemic^4, 23^.

However, it is now known that nosocomial infection during prolonged hospitalization is the primary threat to patients with COVID-19^24^, rather than bacterial co-infection upon hospital admission^9, 25–27^. Evidence from immunocompromised cancer patients suggests that indiscriminate administration of broad-spectrum antibiotics may, counter-intuitively, increase nosocomial bloodstream infection (nBSI) rates by causing gut dominations of resistant microbes that can translocate into the blood^18, 28^. Thus, empiric antimicrobial use, i.e. without direct evidence for a bacterial infection, in patients with severe COVID-19 may be especially pernicious because it may select for antimicrobial resistance and could promote gut translocation-associated nBSI.

The role of the gut microbiome in respiratory viral infections in general^29–31^, and in COVID-19 patients in particular, is only beginning to be understood. Animal models of influenza virus infection have uncovered mechanisms by which the microbiome influences antiviral immunity^32–34^, and in turn, the viral infection was shown to disrupt the intestinal barrier of mice by damaging the gut microbiota^35, 36^. Hence, we hypothesized that gut dysbiosis during COVID-19 may be associated with BSIs. To test this, we first determined whether SARS-CoV-2 infection could directly cause gut dysbiosis independently of hospitalization and treatment. K18-hACE2 mice (*K18-ACE2tg* mice), express human *ACE2* driven by the *cytokeratin-18* promoter (*K18-ACE2tg* mice). Although the overexpression of ACE2 prevents investigation of long term consequences of infection due to potential non-specific disease, which is a major limitation of the model, an advantage of these mice is that they develop severe respiratory disease in a virus dose-dependent manner, partially mirroring what is observed in COVID-19 patients^37–40^. Daily changes in fecal bacterial populations were monitored following intranasal inoculation of mice with a range of doses (10, 100, 1000, and 10000 PFU) of SARS-CoV-2 or mock-treatment (**Fig. 1a**, **Extended Data Fig. S1**). Although we detected viral RNA in the lungs of mice infected with doses as low as 100 PFU (**Extended Data Fig. S1c**), mice inoculated with doses lower than 10000 PFU displayed minimal or no signs of disease (**Extended Data Fig. S1a,b**), and as expected based on this outcome, shifts in their microbiome were inconsistent (**Extended Data Fig. S2**). Thus, we focused on findings from the 10000 PFU inoculum.

**Fig. 1.**
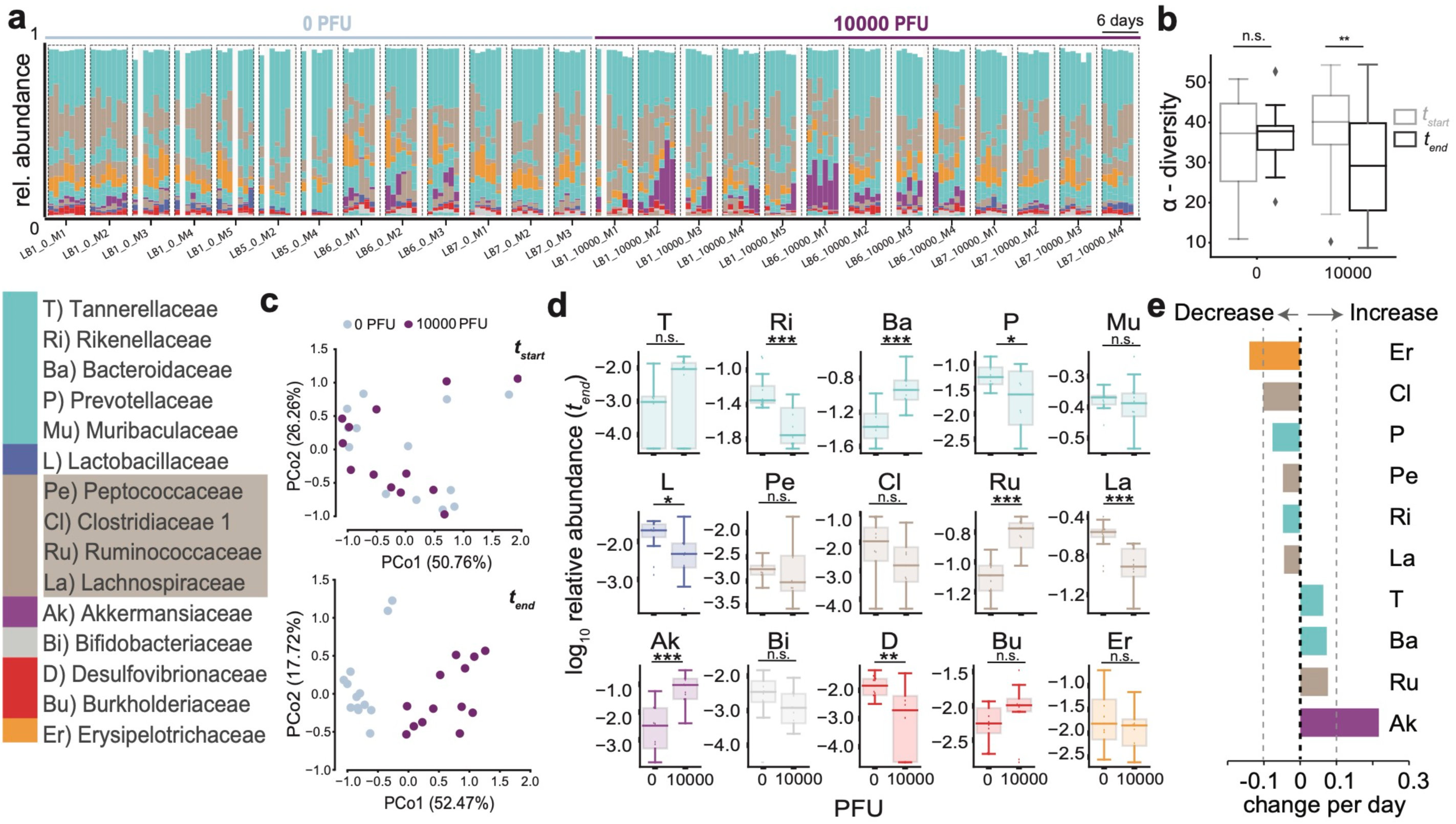
SARS-CoV-2 infection causes gut microbiome alterations in mice. **a** Timelines of fecal microbiota composition measured by 16S rRNA gene sequencing in mice infected with 0 or 10^4^ PFU of SARS-CoV-2 ; time of infection=Day 1. Bars represent the composition of the 15 most abundant bacterial families per sample, blocks of samples correspond to an individual mouse’s time course (x-axis label indicate experiment id, PFU, and mouse id). **b** *α*-diversity (inverse Simpson index) per infection group in the beginning (*_tstart_*) and at the end (*t_end_*) of the experiment (n.s.: non-significant, **: p<0.01, one-tailed, paired t-test). **c** Principal coordinate plot of bacterial compositions in samples from the start (top) and end (bottom) of the experiment. **d** *log*_10_-relative family abundances at the final time point; boxplots show median and interquartile ranges, whiskers extend to 1.5 times max- and min-quartile values, n.s.: not significant; *: p-value < 0.05; **: p-value < 0.01; ***: p-value < 0.001; Wilcoxon rank-sum tests. **e** Regression coefficients of the estimated changes in family abundances per day in mice infected with 10^4^ PFU obtained from linear mixed effects models with varying effects per mouse and per cage (only significant coefficient results shown, abbreviations and colors as per the bacterial family legend).

Mice infected with 10000 PFU displayed weight loss and other signs of disease around day 4 (**Extended Data Fig. S1a,b, S2e,f**), alongside microbiome changes characterized by a significant loss of alpha diversity (inverse Simpson index, **Fig. 1b**) corresponding to shifts in the bacterial community composition (**Fig. 1c,d**). We performed time series analyses on bacterial family abundances, contrasting their trajectories in infected (10000 PFU) and uninfected mice. This revealed that the strongest shift over time in infected mice was characterized by significant increases of Akkermansiaceae (p<0.0002, **Fig. 1d**). Ranking all bacterial family trajectories by their estimated changes over time in infected mice showed that this increase in Akkermansiaceae was accompanied by significant losses of Clostridiaceae 1, a family of obligate anaerobe bacteria, and of Erysipelotrichiaceae (**Fig. 1e**). These results demonstrated that SARS-CoV-2 infection leads to gut microbiome dysbiosis over time in a mouse model.

We then determined if this dysbiosis was also associated with intestinal defects that could enable translocation of bacteria into the blood. While several of the infected mice displayed signs of barrier dysfunction the observed differences in plasma concentrations of Fluorescein isothiocyanate (FITC)-dextran following its administration by gavage, or other markers of intestinal barrier permeability, fatty acid-binding protein (iFABP), Lipopolysaccharide-binding protein (LBP), and citrulline did not reach significance (**Extended Data Fig. S3a,b**). The reduced colon lengths as well as reductions in the villus lengths in the duodenum or ileum, i.e. markers of overt inflammation, that we observed were also non-significant compared with control mice (**Extended Data Fig. S3c,d**). However, infected mice that had incurred the most severe microbiome injury in the form of diversity loss showed the most evidence of gut permeability–the highest FITC-dextran concentrations in the blood of mice detected across all samples came from the two out of the four mice with the most extreme dysbiosis and highest levels of Akkermansiaceae, a family of mucin-degrading bacterial species (**Extended Data Fig. S4)**.

Interestingly, we also detected a significant increase in the number of mucus-producing goblet cells and a decrease in the number of Paneth cells in the ileum (but not in the duodenum) of infected mice (**Fig. 2a,c and Extended Data Fig. S3e**). The decrease in Paneth cells was accompanied by structural abnormalities, most notably deformed or misplaced granules (**Fig. 2b**). These morphological abnormalities in Paneth cells were reminiscent of observations in the ileum of patients with inflammatory bowel disease (IBD) as well as in a virally-triggered animal model of IBD, where such structures were indicative of defects in packaging and secretion of the granule protein lysozyme^41–43^. Thus, to quantify the Paneth cell granule defect, we performed lysozyme immunofluorescence and found a significant increase in the proportion of Paneth cells displaying abnormal staining patterns compared with the controls (**Fig. 2b,c**). We then investigated if these physiological defects were associated with dysbiosis in the microbiome. The most severely sick mice also had the most striking shifts in their microbiome composition and the lowest microbiota diversity at the end of the experiment (**Extended Data Fig. S4a,b**). To associate the observed physiological defects with microbiome dysbiosis, we plotted the numbers of goblet cells per crypt-villus unit and Paneth cells per crypt, as well as the percentage of abnormal Paneth cells against bacterial alpha diversity and the log10-relative abundance of Akkermansiaceae (**Fig. 2d,e**). Goblet cell counts per crypt-villus unit were negatively correlated with alpha diversity, and, conversely, positively correlated with Akkermansiaceae. While no statistically significant association was found between diversity, Akkermansiaceae abundance and Paneth cell counts per crypt, we observed a striking positive correlation between the percentage of abnormal Paneth cells and Akkermansiaceae, and a corresponding negative correlation with diversity. We were unable to reliably detect viral RNA in intestinal samples (**Extended Data Fig. S1c**), raising the possibility that systemic immune responses rather than direct cytotoxicity from local viral infection mediate these changes. Altogether, these results show that the gut microbiome dysbiosis observed in K18-hACE2 mice infected with a high dose of SARS-CoV-2 are associated with alterations in key epithelial cells, and signs of barrier permeability in the mice displaying the greatest disruption in microbiome diversity.

**Fig. 2.**
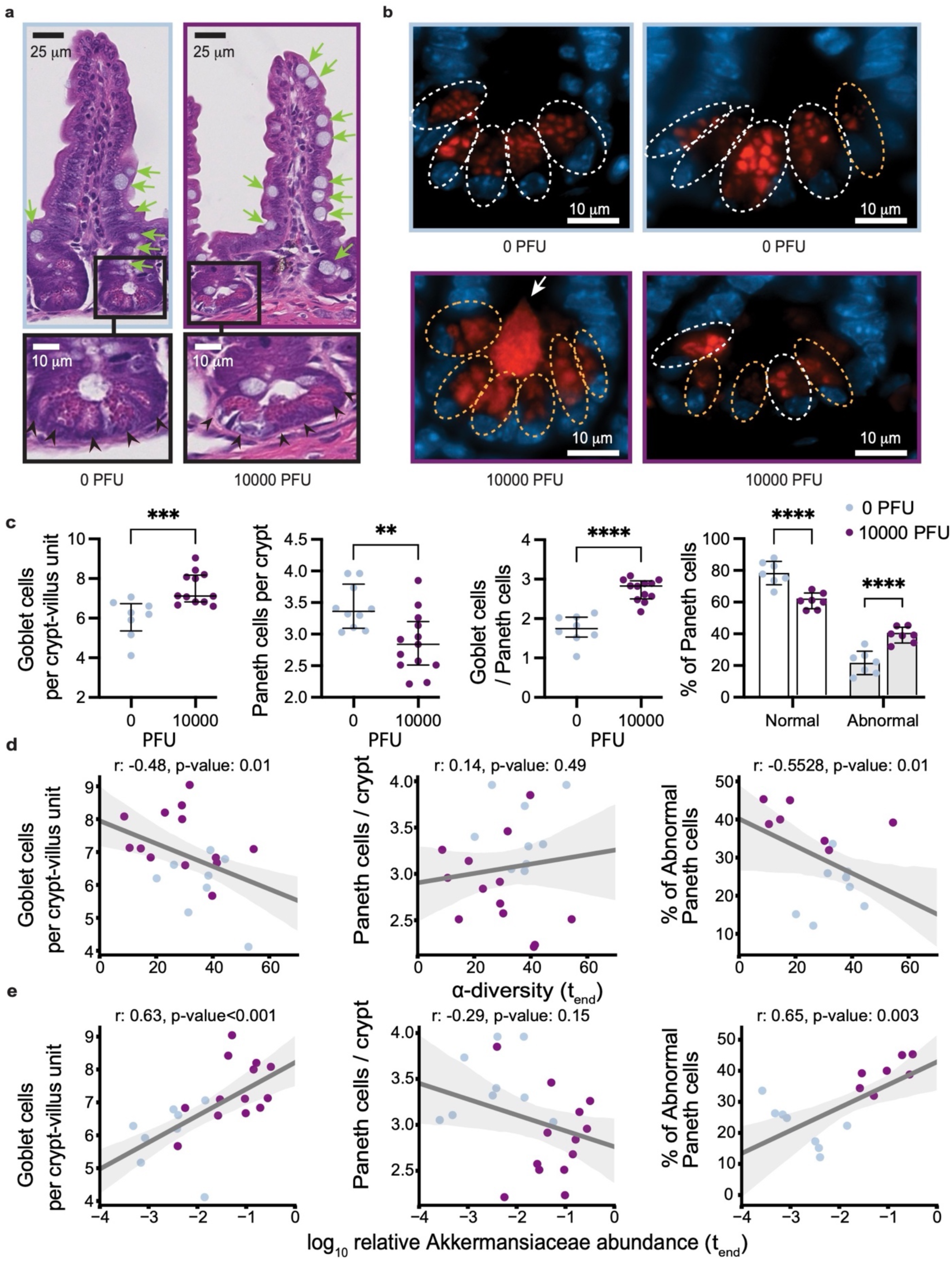
SARS-CoV-2 infection causes abnormalities in the gut epithelium of mice. **a.** Representative H&E-stained section of the ileum depicting crypt-villus axes from K18-hACE2 mice on day 5-6 post intranasal inoculation with 10000 PFU SARS-CoV-2 or mock treatment. Green arrows indicate goblet cells, scale bars correspond to 25μm. Bottom panels show high magnification images of the indicated crypt with black arrowheads pointing at Paneth cells, scale bars correspond to 10μm. **b.** Representative anti-lysozyme immunofluorescence images of the ileal crypt (two images per group). White and orange doted circles delineate normal and abnormal Paneth cells, respectively. Abnormality is characterized by distorted, depleted, or diffuse lysozyme distribution patterns in Paneth cells. Lysozyme = red, DAPI = blue, scale bars correspond to 10 m. **c.** Quantification of goblet cell number per villus (left), Paneth cells per crypt (middle) based on H&E staining, and frequency of normal versus abnormal Paneth cell lysozyme distribution pattern based on the immunofluorescence staining as depicted in b. Dots represent the mean cell number per crypt-villus unit in each mouse, 50 units were counted per mouse. Results were pooled from 3 independent experiments with n=3-5 mice per group for each experiment. Boxplots indicate median and interquartile ranges (ns=non-significant, p>0.05; **, p<0.01; ***, p<0.001; ****, p<0.0001 Mann-Whitney U-test). **d.** Correlation of Goblet cell number per villus (left, Pearson correlation r=-0.48, p=0.015), Paneth cells per crypt (middle, r=0.14, p-value=0.483) and frequency of abnormal Paneth cell lysozyme distribution pattern (right, r=-0.5528, p=0.014) for the mice shown in c with α-diversity (inverse Simpson) of the gut microbiome measured at the last day before sacrifice. **e.** Correlation of Goblet cell number per villus (left, r=0.63, p<0.001), Paneth cells per crypt (middle, r=-0.29, p=0.149) and frequency of abnormal Paneth cell lysozyme distribution pattern (right, r=0.65, p-value=0.003) for the mice shown in c with log_10_-relative abundances of *Akkermansia* in fecal samples from the last day before sacrifice; lines: univariate linear regression, shaded region: 95% CI.

To investigate the microbiome in COVID-19 patients, we profiled the bacterial composition of the fecal microbiome in 130 samples (**Fig. 3a**) obtained from SARS-CoV-2 infected patients treated at NYU Langone Health (NYU, 67 samples from 60 patients) and Yale New Haven Hospital (YALE, 63 samples from 36 patients, **Supplementary Table 1**). Analysis of metagenomic data obtained from sequencing of the 16S rRNA genes revealed a wide range of bacterial community diversities, as measured by the inverse Simpson index, in samples from both centers (NYU: [1.0, 32.3], YALE: [1.5, 29.3], **Fig. 3b**); on average, samples from NYU were less diverse (-4, p<0.01, two-tailed T-test, **Fig. 3c**), and as reported previously, samples from patients admitted to the ICU had reduced diversity (-3.9, p<0.05, two-tailed T-test, **Extended Data Fig. S5a**). However, the composition in samples between the two centers did not show systematic compositional differences (**Fig. 3d,e,f**). On average, in both centers, members of the phyla Firmicutes and Bacteroidetes represented the most abundant bacteria, followed by Proteobacteria (**Fig. 3d**). The wide range of bacterial diversities was reflected in the high variability of bacterial compositions across samples (**Fig. 3e,f**). In samples from both centers, microbiome dominations, defined as a community where a single genus reached more than 50% of the population, were observed frequently (NYU: 21 samples, YALE: 12 samples), representing states of severe microbiome injury in COVID-19 patients (**Fig. 3g, Extended Data Fig. S5a,b**). Strikingly, samples associated with a BSI, defined here as a positive clinical blood culture test result, had strongly reduced bacterial α-diversities (mean difference: -5.2, CI_BEST_[-8.2, -2.2], **Fig. 3h**).

**Fig. 3.**
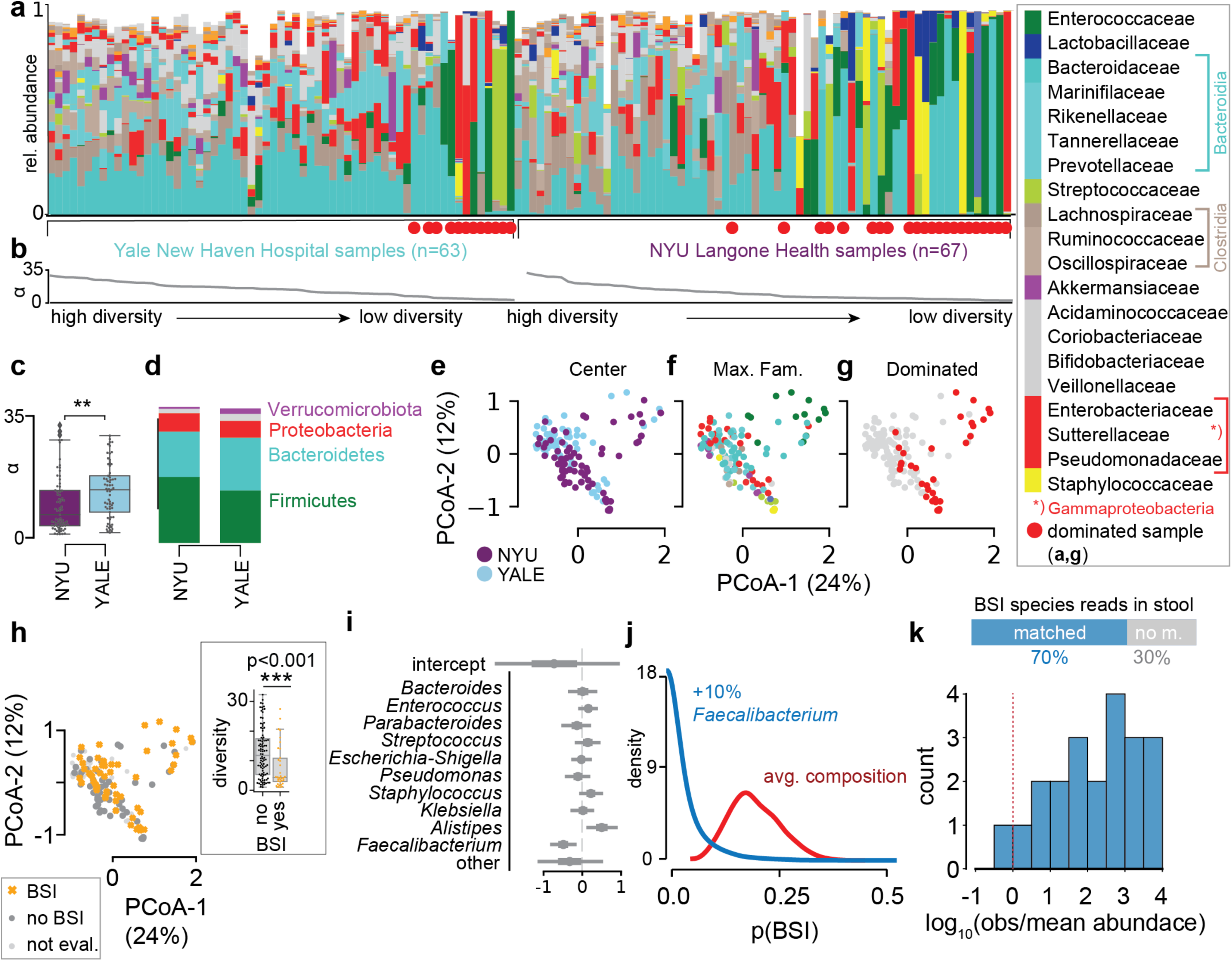
The dysbiotic gut microbiome in COVID-19 in patients from NYU Langone Health (n=60) and Yale New Haven Hospital (n=36) is associated with secondary bloodstream infections. **a.** Bacterial family composition in stool samples (Yale, n = 63 samples; NYU, n = 67) identified by 16S rRNA gene sequencing; bars represent the relative abundances of bacterial families; red circles indicate samples with single taxa *>*50%. Samples are sorted by center and bacterial *α*-diversity (inverse Simpson index, **b**). c *α*-diversity in samples from NYU Langone Health and Yale New Haven Hospital; **p<0.01, two-sided T-test. **d** Average phylum level composition per center. **e-g** Principal coordinate plots of all samples shown in **a**, labeled by center (**e**), most abundant bacterial family (**f**) and domination status of the sample (**g**), and BSI status; inset: boxplot of inverse Simpson index diversity by BSI (**h**). **i** Coefficients from a Bayesian logistic regression with most abundant bacterial genera as predictors of BSI status. **j** Counterfactual posterior predictions of BSI risk based on bacterial composition contrasting the predicted risk of the average composition across all samples (red) with the risk predicted from a composition where *Faecalibacterium* was increased by 10% (blue). **k** shotgun metagenomic reads matched the species identified in clinical blood cultures in 70% of all investigated cases; the histogram shows the distribution of log_10_-ratios of relative abundances of matched species in corresponding stool samples to their corresponding mean abundances across all samples.

The lower diversity associated with samples from 25 patients (15 NYU, 10 Yale) with BSIs led us to investigate their bacterial taxon compositions and the potential that gut dysbiosis was associated with BSI events. Importantly, BSI patients had received antibiotic treatments during hospitalization (**Extended Data Fig. S6, Supplementary Table 2**), which could exacerbate COVID-19 induced shifts in microbiota populations^16, 17, 20^, and may indeed be administered in response to a suspected or confirmed BSI. We noted that most BSI patients received antibiotics prior to their BSI, with 6 out of 25 patients receiving antibiotics only after detection of BSI. Principal coordinate analysis of all stool samples indicated that the BSI-associated samples spanned a broad range of compositions (**Fig. 3h**). To identify bacterial abundance patterns that consistently distinguished BSI from non-BSI-associated samples, we performed a Bayesian logistic regression. The model estimated the association of the 10 most abundant bacterial genera with BSI cases, i.e. it identified enrichment or depletion of bacterial genera in BSI associated samples (**Fig. 3i**). This analysis revealed that the genus *Faecalibacterium* was negatively associated with BSI (OR: -0.5, CI:[-0.86, -0.15]), which was also observed when we included microbiome domination as an additional factor in the model (**Extended Data Fig. S7a**). However, our analysis also included stool samples that were taken only after a positive blood culture was obtained, calling into question the plausibility of gut translocation; a complementary analysis only using stool samples obtained prior or on the same day of a positive blood culture also identified *Faecalibacterium* as most negatively associated with BSI (**Extended Data Fig. S7b**). Furthermore, a higher-resolution analysis using amplicon sequencing variant (ASV) relative abundances as predictors of BSI (**Extended Data Fig. S7c,d**), identified an ASV of the *Faecalibacterium* genus as most negatively associated with BSI, in agreement with our main analysis. *Faecalibacterium* is an immunosupportive, short-chain fatty acid producing genus that is a prominent member of the human gut microbiome^44–46^, and its reduction is associated with disruption to intestinal barrier function^47, 48^, perhaps via ecological network effects^48^.

To evaluate the effect size of the association between *Faecalibacterium* and BSIs, we performed a counterfactual posterior predictive check. Using the average genus composition found across all samples, we first computed the distribution of predicted BSI risks (**Fig. 3j**), and compared this risk distribution with a hypothetical bacterial composition which increased *Faecalibacterium* by 10% points. The predicted risk distributions associated with these two compositions differed strongly (mean difference 15%, CI: [1%, 32%], **Fig. 3j**). Domination states of the microbiome increase the risk for BSIs in immunocompromised cancer patients^18^; such dominations imply high relative abundances of single taxa, and therefore a low diversity. Consistent with this, *Faecalibacterium* abundance was positively correlated with diversity (R: 0.55, p<10^−10^, **Extended Data Fig. S8**) in our data set and as reported previously^44^.

We therefore next investigated a direct association between the bacteria populating the gut microbiome and the organisms identified in the blood of patients. Visualizing the bacterial composition in stool samples from patients alongside the BSI microorganisms (**Extended Data Fig. S9a)** suggested a correspondence with the respective taxa identified in the blood: high abundances of the BSI-causing microbes were found in corresponding stool samples. A rank abundance analysis matching the organisms identified in clinical blood cultures to the composition of bacteria in corresponding stool samples indicated enrichment of taxa belonging to the same bacterial orders as BSI causing organisms (**Extended Data Fig. S9b**), suggesting translocation of bacteria from the gut into the blood stream.

To further investigate evidence for translocation of gut bacteria into the blood, we next performed shotgun metagenomic sequencing on a subset of BSI-associated samples with sufficient remaining material in order to match the organism identified in clinical blood cultures at the species level with reads obtained from stool samples (**Fig. 3k, Supplementary Table 3**). In four cases of positive blood cultures of *Staphylococcus* species, no reads matching the clinically identified species were detected (**Supplementary Table 3**). This may explain why the rank analysis suggested that Staphylococcales were not generally enriched in BSIs by *Staphylococcus* (**Extended Data Fig. S9a,b**). In all investigated cases of positive blood cultures by other organisms, the species identified in clinical blood cultures had corresponding reads in the stool samples. Furthermore, the relative abundances of matched species tended to be larger than the average abundances of matched species across all samples (**Supplementary Table 3**). Consistent with this, in one case of a *S. aureus* BSI where corresponding stool relative abundances of *Staphylococcus* were low, reads from shotgun sequencing did not match the genomes of isolates obtained from the same patient better than *S. aureus* genomes from other isolates (**Extended Data Fig. S9d**). Strikingly, shotgun metagenomic reads matched the genome of isolates well in another case where relative abundances of *Staphylococcus* were enriched in the stool (**Extended Data Fig. S9c**), providing evidence that here, the same strains were found in stool and blood of the same patient.

Collectively, these results reveal an unappreciated link between SARS-CoV-2 infection, gut microbiome dysbiosis, and a severe complication of COVID-19, BSIs. The loss of diversity and immunosupportive *Faecalibacterium* in patients with BSIs mirrored a similar loss of diversity in the most severely sick mice deliberately infected with SARS-CoV-2, and as observed by other labs and other model systems^49–51^. Notably, a recent study reproduced these changes in the microbiome in an antibiotics-naïve cohort^52^, suggesting that the viral infection causes gut dysbiosis, either through gastrointestinal infection^53–57^ or through a systemic inflammatory response^2, 4^. Furthermore, the pronounced increase in Akkermansiaceae in mice was also observed in our patient samples, and has been reported previously in patients and in K18-hACE2 mice^49, 58^. However, the dysbiosis in patients with COVID-19 exceeded the microbiota shifts observed in the mouse experiments, including microbiome dominations by single taxa, which was not seen in the mouse experiments. It is possible that in our experiment, mice were sacrificed before perturbations to the gut microbial populations reached a maximum. hACE2 knock-in mice, which display reduced disease^37^, were not tested in the scope of this study but could provide additional insights in the future. However, it is also plausible that the frequently administered antibiotic treatments that hospitalized COVID-19 patients receive exacerbated SARS-CoV-2 induced microbiome perturbations. Additionally, unlike the controlled environment experienced by laboratory mice, hospitalized patients are uniquely exposed to antimicrobial-resistant infectious agents present on surfaces and shed by other patients.

Despite these limitations of the mouse model, we observed that SARS-CoV-2 infection led to alteration of intestinal epithelial cells with established roles in intestinal homeostasis and gastrointestinal disease^59, 60^. Microbiome ecosystem shifts are likely both cause and consequence of these epithelial cell alterations, since epithelial secretions are predicted to affect overall community structure disproportionately strongly^61, 62^. For example, disruption of Paneth cell-derived antimicrobials including lysozyme are sufficient to impact microbiome composition^63–65^, and, conversely, *Akkermansia*, which was increased in infected mice, can have epithelium remodeling properties^66^.

Our observation that the type of bacteria that entered the bloodstream was enriched in the associated stool samples is a well characterized phenomenon in cancer patients^18^, especially during chemotherapy induced leukocytopenia when patients are severely immunocompromised^16, 44^. COVID-19 patients are also immunocompromised and frequently incur lymphopenia, rendering them susceptible to secondary infections^67^. Our data suggests dynamics in COVID-19 patients may be similar to those observed in cancer patients: BSI-causing organisms may translocate from the gut into the blood, potentially due to loss of gut barrier integrity, through tissue damage downstream of antiviral immunity rather than chemotherapy. Consistent with this possibility, soluble immune mediators such as TNF*α* and interferons produced during viral infections, including SARS-CoV-2, damage the intestinal epithelium to disrupt the gut barrier, especially when the inflammatory response is sustained as observed in patient with severe COVID-19^43, 68, 69^. Indeed, blood plasma in severely sick COVID-19 patients are enriched for markers of disrupted barrier integrity and higher levels of inflammation markers^70^, suggesting microbial translocation. Our data supports this model with direct evidence because we were able to match sequencing reads from stool samples to genomes of species detected in the blood of patients.

One limitation of our data is temporal ordering of samples. Occasionally stool samples were collected after observation of BSI, and this mismatch in temporal ordering is counter intuitive for gut-to-blood translocation and a causal interpretation of our associations. However, the reverse direction, that blood infection populates and changes the gut community, is unlikely for the organisms identified in the blood, and if our associations were not causal, we would expect no match between BSI organisms and stool compositions.

Taken together, our findings support a scenario in which gut-to-blood translocation of microorganisms following microbiome dysbiosis, a known issue for chronic conditions such as cancer, leads to dangerous BSIs during COVID-19. We suggest that investigating the underlying mechanism behind our observations will inform the judicious application of antibiotics and immunosuppressives in patients with respiratory viral infections and increase our resilience to pandemics.

## Materials and Methods

### Bioethics statement

The collection of COVID-19 human biospecimens for research has been approved by the NYUSOM Institutional Review Board under il8-01121 Inflammatory Bowel Disease and Enteric Infection at NYU Langone Health. The data presented in this study were also approved by Yale Human Research Protection Program Institutional Review Boards (FWA00002571, protocol ID 2000027690). Informed consent was obtained from all enrolled patients.

### Mouse experiments

#### Cells & virus

Vero E6 (CRL-1586; American Type Culture Collection) were cultured Dulbecco’s Modified Eagle’s Medium (DMEM, Corning) supplemented with 10% fetal bovine serum (FBS, Atlanta Biologics) and 1% nonessential amino acids (NEAA, Corning). SARS-CoV-2, isolate USA-WA1/2020 19 (BEI resources #NR52281), a gift from Dr. Mark Mulligan at the NYU Langone Vaccine Center was amplified once in Vero E6cells. All experiments with SARS-CoV-2 were conducted in the NYU Grossman School of Medicine ABSL3 facility in accordance with its Biosafety Manual and Standard Operating Procedures, by personnel equipped with powered air-purifying respirators.

#### Mice

Heterozygous K18-hACE2 C57BL/6J mice (strain: 2B6.Cg-Tg(K18-ACE2)2Prlmn/J) were obtained from The Jackson Laboratory. Several were paired with C57BL/6J mice to generate additional heterozygous mice for subsequent experiments and the remaining were used to perform initial experiments. Animals from the same breeder pool (i.e., littermates) were randomly assigned and housed in cages according to the experimental groups and fed standard chow diets. Cage bedding was mixed prior to infection in a subset of experiments to further reduce possible cage effect. All animal studies were performed according to protocols approved by the NYU School of Medicine Institutional Animal Care and Use Committee (IACUC n°170209 and 180802) and in the ABSL3 facility of NYU Grossman School of Medicine (New York, NY), in accordance with its Biosafety Manual and Standard Operating Procedures. 12-week-old or 24-week-old K18-hACE2 males were administered either 10-10000 PFU SARS-CoV-2 diluted in 50µL PBS (Corning) or 50µL PBS (non-infected, 0) via intranasal administration under xylazine-ketamine anesthesia (AnaSedR AKORN Animal Health, KetathesiaTM Henry Schein Inc). Viral titer in the inoculum was verified by plaque assay in Vero E6 cells. Following infection, mice were monitored daily for weight loss, temperature loss and signs of disease. A disease score was calculated as the sum of scores obtained for each of the following criteria: ruffled fur (no= 0, yes=1), hunched back (no=0, slightly=1, exacerbated=2), heavy breathing (no=0, yes=1), altered mobility (no=1, reduced activity=1, no mobility=2). Stool samples were collected and stored at -80°C.

#### Measurement of viral load

Whole lungs and 1cm of proximal duodenum, terminal ileum and proximal colon were collected five-to seven days after infection. Intestinal pieces were wash with PBS and all organs were transferred in Eppendorf tubes containing 500μl of PBS and a 5mm stainless steel bead (Qiagen) and h C57BL/6J mice omogenized using the Qiagen TissueLyser II. Homogenates were cleared for 5 min at 5,000 × g, and the viral supernatant or nasal wash was diluted 4X in TRIzol reagent (Invitrogen) and frozen at -80°C for titration by qRT-PCR. RNA was extracted from the TRIzol homogenates using chloroform separation and isopropanol precipitation, followed by additional purification using RNeasy spin columns with DNase treatment according to the manufacturer’s instructions (Rneasy Mini Kit; RNAse-Free DNase Set; QIAGEN). RNA was reverse-transcribed using the High-Capacity cDNA Reverse Transcription Kit (Applied Biosystems). qPCR was performed using Applied Biosystems TaqMan RNA-to-CT One-Step Kit (Fisher-Scientific), 500nM of the primers (Fwd 5’-ATGCTGCAATCGTGCTACAA-3’, Rev 5’-GACTGCCGCCTCTGCTC-3’) and 100nM of the N probe (5’-/56-FAM/TCAAGGAAC/ZEN/AACATTGCCAA/3IABkFQ/-3’). Serial dilutions of *in-vitro* transcribed RNA of the SARS-CoV-2 Nucleoprotein (generated as previously described^71^) were used to generate a standard curve and calculate copy numbers per ug of RNA in the samples.

#### Microscopy

5cm of proximal duodenum, distal ileum and entire colon were flushed with 10% acetate buffered formalin (Fisher scientific), cut open along the length, pinned on black wax and fixed with formalin for 72hrs at RT. 2 cm strips of intestinal tissues were embedded in low melting point agarose (Promega) to enrich for well-oriented crypt-villus units. Paraffin embedding, sectioning, and staining were performed by the NYU Experimental Pathology Research Laboratory. 5um sections were stained with hematoxylin and eosin (H&E) and imaged using brightfield wholeslide scanning. Lysozyme staining was performed using anti-lysozyme (ab108508, Abcam) and DAPI immunostaining and analyzed using a Zeiss AxioObserver.Z1 with Axiocam 503 Mono operated with Zen Blue software. 50 small intestinal villi per mouse were measured for villi length. Goblets cell were quantified from 50 villus-crypt units (one villus + half of the 2 surrounding crypts) per mouse. Paneth cells numbers and lysozyme staining patterns were quantified from 50 crypts per mouse. Previously defined criteria were used to quantify the proportion of Paneth cells displaying abnormal lysozyme staining^43^. Mean values were calculated for each mouse and used as individual data points.

#### Measurement of intestinal permeability

Mice were fasted for 4hrs before oral gavage with 200uL of fluorescein isothiocyanate (FITC)-dextran (3-5 kDa, Sigma-Aldrich) dissolved in sterile PBS (60mg/ml). After 4 hrs, mice were euthanized and blood was collected by cardiac puncture. FITC-dextran in plasma was quantified using a plate reader (excitation, 485 nm; emission, 530 nm). Citrulline, intestinal fatty acid-binding protein, and lipopolysaccharide (LPS)-binding protein were quantified in the plasma by enzyme-linked immunosorbent assay (ELISA) according to the manufacturer’s instructions (MyBioSource, CA).

#### Time series analyses of bacterial family abundances

We log_10_-transformed bacterial relative abundances adding a pseudo count to fill zeros (2*10^-6^, as done before). We then analyzed the time series with the following model that included fixed effects for the intercepts and slopes of the treatment (i.e. indicator variables for uninfected (0 PFU), and infected (10000PFU), and random effects per cage and per mice to account for cage effects and repeated measurements from the same individual mouse, respectively. The model was implemented in the R programming language using the lmer function of the lme4 library with the following model formula:

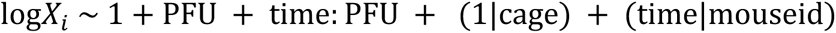

### Human study

#### Study population and data collection

This study involved 96 patients with laboratory-confirmed SARS-CoV-2 infection. SARS-CoV-2 infection was confirmed by a positive result of real-time reverse transcriptase-polymerase chain reaction assay on a nasopharyngeal swab. 60 patients were seen at NYU Langone Health, New York, between January 29, 2020 – July 2, 2020. In order to be eligible for inclusion in the study, stool specimens needed to be from individuals >18 years of age. Data including demographic information, clinical outcomes, and laboratory results were extracted from the electronic medical records in the NYU Langone Health clinical management system. Blood and stool samples were collected by hospital staff. OmnigeneGut kits were used on collected stool. In parallel, 36 patients were admitted to YNHH with COVID-19 between 18 March 2020 and 27 May 2020 as part of the YALE IMPACT cohort described at length elsewhere^2^. Briefly, participants were enrolled after providing informed consent and paired blood and stool samples were collected longitudinally where feasible for duration of hospital admission. No statistical methods were used to predetermine sample size for this cohort. Demographic information of patients was aggregated through a systematic and retrospective review of the EHR and was used to construct **Supplementary Table 1**. Symptom onset and etiology were recorded through standardized interviews with patients or patient surrogates upon enrolment in our study, or alternatively through manual EHR review if no interview was possible owing to clinical status at enrolment. The clinical data were collected using EPIC EHR and REDCap 9.3.6 software. At the time of sample acquisition and processing, investigators were blinded to patient clinical status.

#### DNA extraction and bacterial 16S rRNA sequencing

For bacterial DNA extraction 700µL of SL1 lysis buffer (NucleoSpin Soil kit, Macherey-Nagel) was added to the stool samples and tubes were heated at 95°C for 2h to inactivate SARS-CoV-2. Samples were then homogenized using the FastPrep-24TM instrument (MP Biomedicals) and extraction was pursued using the NucleoSpin Soil kit according to the manufacturer’s instructions. DNA concentration was assessed using a NanoDrop spectrophotometer. Samples with too low DNA concentration were excluded. DNA from human samples was extracted with PowerSoil Pro (Qiagen) on the QiaCube HT (Qiagen), using Powerbead Pro (Qiagen) plates with 0.5mm and 0.1mm ceramic beads. For mouse samples, the variable region 4 (V4) of the 16S rRNA gene was amplified by PCR using primers containing adapters for MiSeq sequencing and single-index barcodes. All PCR products were analyzed with the Agilent TapeStation for quality control and then pooled equimolar and sequenced directly in the Illumina MiSeq platform using the 2x250 bp protocol. Human samples were prepared with a protocol derived from ^72^, using KAPA HiFi Polymerase to amplify the V4 region of the 16S rRNA gene. Libraries were sequenced on an Illumina MiSeq using paired-end 2x250 reads and the MiSeq Reagent Kitv2.

#### Bioinformatic processing and taxonomic assignment

Amplicon sequence variants (ASVs) were generated via dada2 v1.16.0 using post-QC FASTQ files. Within the workflow, the paired FASTQ reads were trimmed, and then filtered to remove reads containing Ns, or with maximum expected errors >= 2. The dada2 learn error rate model was used to estimate the error profile prior to using the core dada2 algorithm for inferring the sample composition. Forward and reverse reads were merged by overlapping sequence, and chimeras were removed before taxonomic assignment. ASV taxonomy was assigned up to genus level using the SILVAv.138 database with the method described in ^73^ and a minimum boostrapping support of 50%. Species-level taxonomy was assigned to ASVs only with 100% identity and unambiguous matching to the reference.

#### Shotgun metagenomic sequencing

DNA was quantified with Qiant-iT Picogreen dsDNA Assay (Invitrogen). Libraries were prepared with a procedure adapted from the Nextera Library Prep kit (Illumina), and sequenced on an Illumina NovaSeq using paired-end 2x150 reads (Illumina) aiming for 100M read depth. DNA sequences were filtered for low quality (Q-Score < 30) and length (< 50), and adapter sequences were trimmed using cutadapt. Fastq files were converted a single fasta using shi7. Sequences were trimmed to a maximum length of 100 bp prior to alignment. DNA sequences were taxonomically classified using the MetaPhlAn2 analysis tool (http://huttenhower.sph.harvard.edu/metaphlan2). MetaPhlAn2 maps reads to clade-specific marker genes identified from ∼17,000 reference genomes and estimates clade abundance within a sample from these mappings.

#### Mapping shotgun reads to whole genome sequences of clinical isolates

Quality-controlled reads were re-classified using Kraken2 (Minikraken2 v2 database, available on https://ccb.jhu.edu/software/kraken2/index.shtml). Reads that were classified by Kraken2 as *Staphylococcus aureus* (or a strain thereof) were further mapped using Bowtie2 separately to each of a collection of *Staphylococcus aureus* isolates. The collection was composed of all NCBI RefSeq assemblies as of 11/17/2021, in addition to *Staphylococcus aureus* isolates that were isolated from our subjects. Bowtie2 mapped reads were then further filtered, keeping only reads that mapped without mismatches. A neighbor-joining (NJ) tree was produced from this collection of genomes using Snippy (https://github.com/tseemann/snippy).

### Compositional analyses

#### α-Diversity

We calculated the inverse Simpson (*IVS*) index from relative ASV abundances (*p*) with *N* ASVs in a given sample, 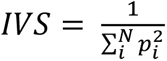

#### Principal Coordinate Analyses

Bray-Curtis distances were calculated from the filtered ASV table using QIIME 1.9.1 and principal components of the resulting distance matrix were calculated using the scikit-learn package for the Python programming language, used to embed sample compositions in the first two principal coordinates.

#### Average compositions and manipulation of compositions

To describe the average composition of a set of samples we calculated the central tendency of a compositional sample ^74^. For counter factual statistical analyses that require changes to a composition, e.g. an increase in a specific taxon, we deployed the perturbation operation (⊕), which is the compositional analogue to addition in Euclidean space^74^. A sample *x* containing the original relative taxon abundances is perturbed by a vector *y*,

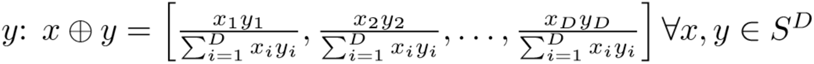

where *S^D^* represents the D-part simplex.

#### Bayesian t-test

To compare diversity measurements between different sample groups, e.g. different clinical status, we performed a Bayesian estimation of group differences (BEST, ^75^), implemented using the pymc3 package for the Python programming language; with priors (∼) and deterministic calculations (=) to assess differences in estimated group means as follows:

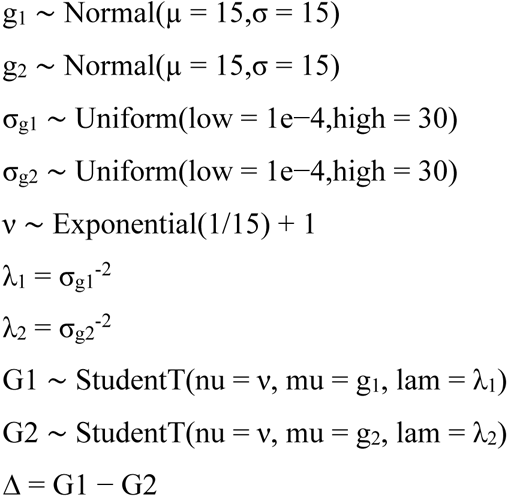

Bayesian inference was performed using “No U-turn sampling”^76^. Highest density intervals (HDI) of the posterior estimation of group differences (Δ) were used to determine statistical certainty (***: 99% HDI >0 or <0, **: 95%HDI, *:90% HDI). The BEST code was implemented following the pymc3 documentation.

#### Cross-validated logistic regression to associate BSI cases with ASV composition

We first removed ASVs with low prevalence (present in fewer than 5% of all samples), and low abundances (maximum observed relative abundance <0.01) leaving 269 ASVs. We then scaled the ASV relative abundances between 0 and 1 (min-max scaling) and performed logistic regressions, relating ASV abundances to BSI status (1: BSI, 0: non-BSI) using the sklearn.linear_model. LogisticRegressionCV module for the Python programming language with an L1 (lasso) penalty, iterating over a range of regularization strengths ([0.01,0.1, 1., 10., 100., 1000.]) using the “liblinear” solver. We retained the inferred ASV association coefficients with non-zero values for each tested regularization strength to visualize the cross-validation path.

#### Bayesian logistic regression

We performed a Bayesian logistic regression to distinguish compositional differences between infection-associated samples and samples from patients without secondary infections. We modeled the infection state of patient sample *i*, *yi* with a Binomial likelihood:

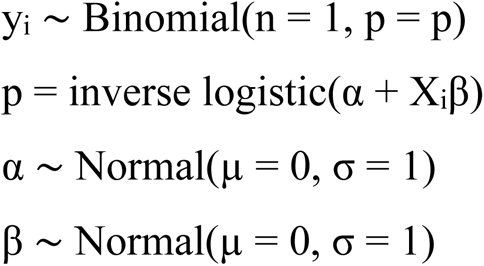

Where prior distributions are indicated by ∼; α is the intercept of the generalized linear model, β is the coefficient vector for the log_10_-relative taxon abundances *X*_i_ in sample *i* or, in some cases, the binary indicator variable for gut microbiome domination.

## Data Availability

The raw sequencing data have been deposited on the Sequencing Reads Archive (SRA), and SRA accession numbers are available for two bioprojects corresponding to the mouse sequencing data PRJNA745367 (**Supplementary Table 4**) and the human stool samples PRJNA746322 (**Supplementary Table 5**).

## Acknowledgments

We thank René Niehus for helpful discussions on the implementation of the various Bayesian analyses. We thank the NYU Langone’s Genome Technology Center, the NYU Langone’s Experimental Pathology Research Laboratory and the NYU Langone’s Microscopy Laboratory supported in part by NYU Langone Health’s Laura and Isaac Perlmutter Cancer Center Support (grant P30CA016087) from the National Cancer Institute Langone and by the NIH S10 OD021747 grant for use of their instruments and technical assistance. We also thank the Office of Science & Research High-Containment Laboratories at NYU Grossman School of Medicine for their support in the completion of this research.

## Yale IMPACT Team

Abeer Obaid, Alice Lu-Culligan, Allison Nelson, Anderson Brito, Angela Nunez, Anjelica Martin, Annie Watkins, Bertie Geng, Chaney Kalinich, Christina Harden, Codruta Todeasa, Cole Jensen, Daniel Kim, David McDonald, Denise Shepard, Edward Courchaine, Elizabeth B. White, Eric Song, Erin Silva, Eriko Kudo, Giuseppe DeIuliis, Harold Rahming, Hong-Jai Park, Irene Matos, Jessica Nouws, Jordan Valdez, Joseph Fauver, Joseph Lim, Kadi-Ann Rose, Kelly Anastasio, Kristina Brower, Laura Glick, Lokesh Sharma, Lorenzo Sewanan, Lynda Knaggs, Maksym Minasyan, Maria Batsu, Mary Petrone, Maxine Kuang, Maura Nakahata, Melissa Campbell, Melissa Linehan, Michael H. Askenase, Michael Simonov, Mikhail Smolgovsky, Nicole Sonnert, Nida Naushad, Pavithra Vijayakumar, Rick Martinello, Rupak Datta, Ryan Handoko, Santos Bermejo, Sarah Prophet, Sean Bickerton, Sofia Velazquez, Tara Alpert, Tyler Rice, William Khoury-Hanold, Xiaohua Peng, Yexin Yang, Yiyun Cao & Yvette Strong

## Author contributions

LBR performed the mouse experiments with help from MGN, AMVJ. CZ performed mouse microbiome analyses with help from LBR, MV and KC. MV, JEA and JS prepared the samples from NYU. MV, JEA prepared the clinical data from NYU with help from JG, EW, BS. JK provided the data from Yale with help from ACM and the IMPACT team, AIK and AI. JS designed and performed the analyses with CZ, and help by GAH and APS. JS and KC designed the research question with support from VJT and BS. JS and KC wrote the manuscript with help by LBR, MV and CZ. All other authors contributed materials, scientific feedback and commented on the manuscript.

## Supporting Information

**Extended Data Fig. S1.**
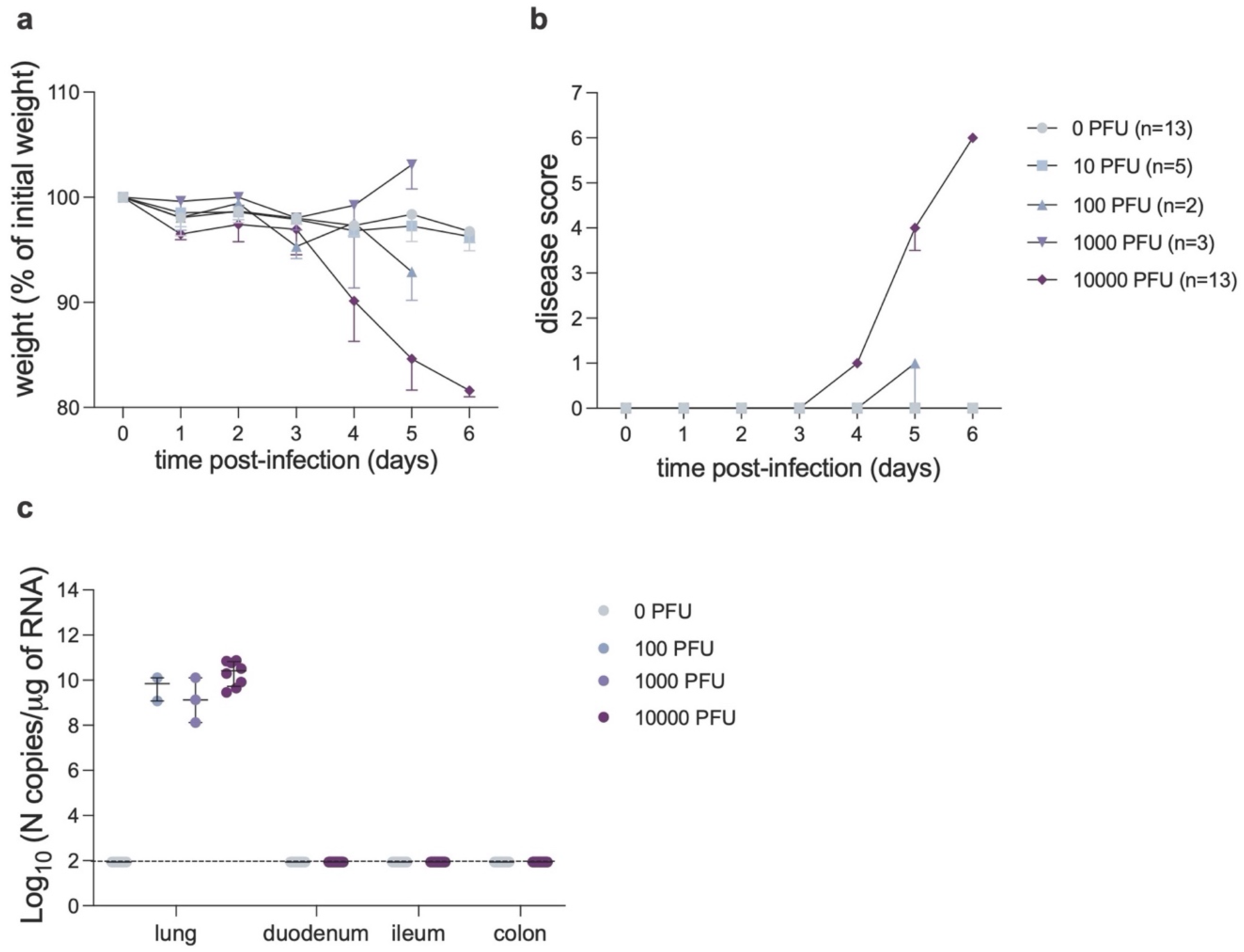
SARS-CoV-2 infection in K18-hACE2 mice. **a-b**. Following inoculation with 0, 10, 100, 1000 or 10000 PFU of SARS-CoV-2 or mock infection, mice were monitored daily for weight loss (a) and signs of disease quantified by a composite score based on ruffled fur, hunched back, heavy breathing and absence of mobility (b). Median and interquartile range determined for each group at each time point are depicted. Results are pooled from 1-3 independent experiments. For each group, the total number of mice is indicated. c. Viral burden in lung or intestinal tissue of K18-hACE2 mice was analyzed at 5-6 days after infection with 100, 1000, 10000 PFU of SARS-CoV-2 or mock infection by qRT-PCR. Dots represent the copy number of N RNA per μg of RNA calculated for each mouse. Results were pooled from 1 (100 and 1000 PFU doses) or 2 (mock and 10000 PFU) independent experiments with n=2-5 mice per group for each experiment. The median and interquartile range are depicted for each experimental group. The dotted line depicts the limit of detection.

**Extended Data Fig. S2.**
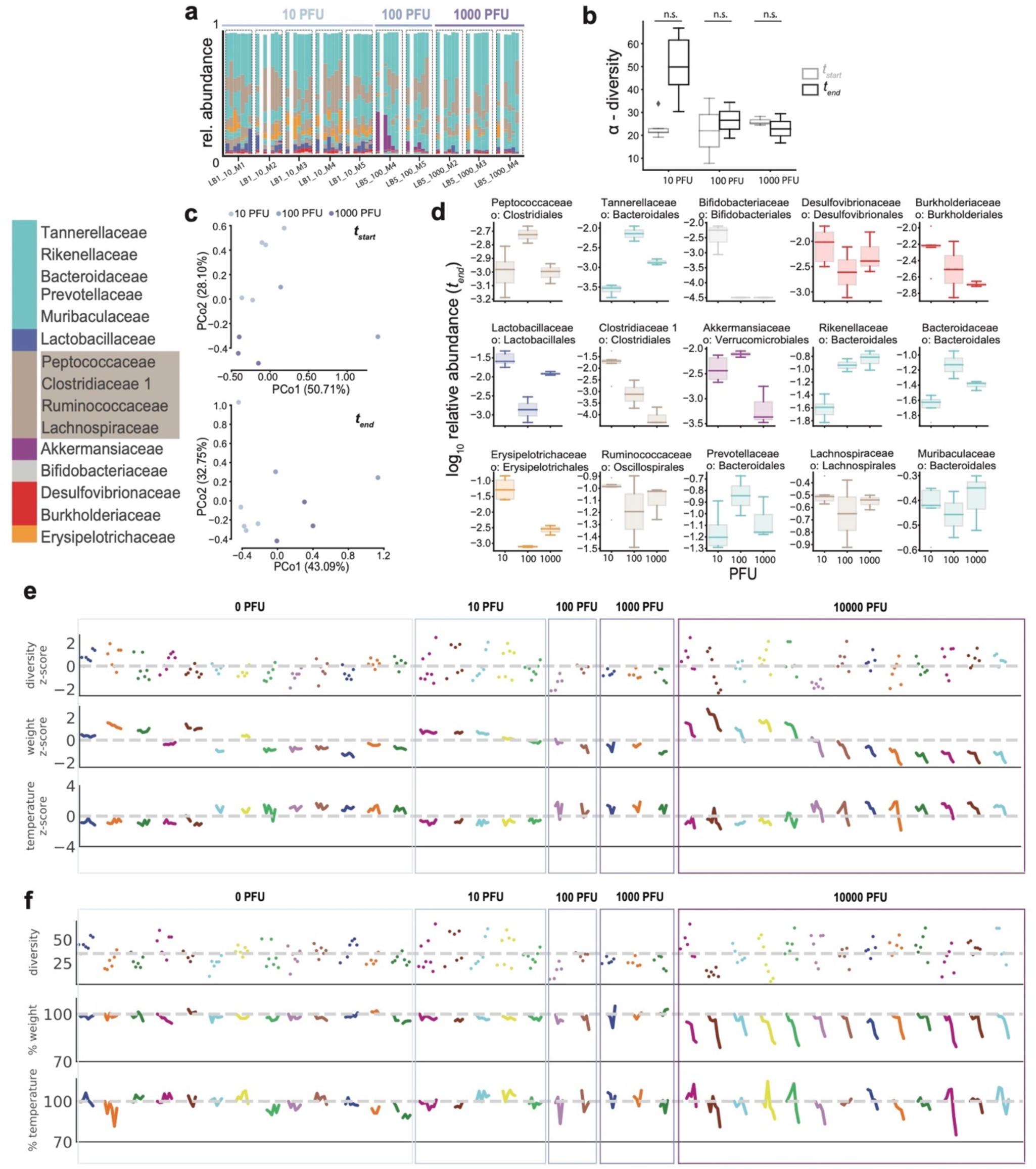
Inconsistent microbiomes dynamics in mice with lower infection doses. **a.** Bars represent bacterial family compositions in stool samples collected from mice over time, mouse time courses grouped as indicated by boxes. **b** bacterial alpha diversity in first (t_start_) and last (t_end_) samples collected. **c** principal coordinate plots of bacterial compositions in first and last samples colored by infection dose (in PFU). **d** bacterial family abundances by infection dose at the final sample collected. **E** diversity, weight and temperature z-scores (calculated from all data points) over time per mouse as shown in a and Fig. 1. **F** untransformed diversity, weights and temperatures relative to the beginning of the experiment.

**Extended Data Fig. S3.**
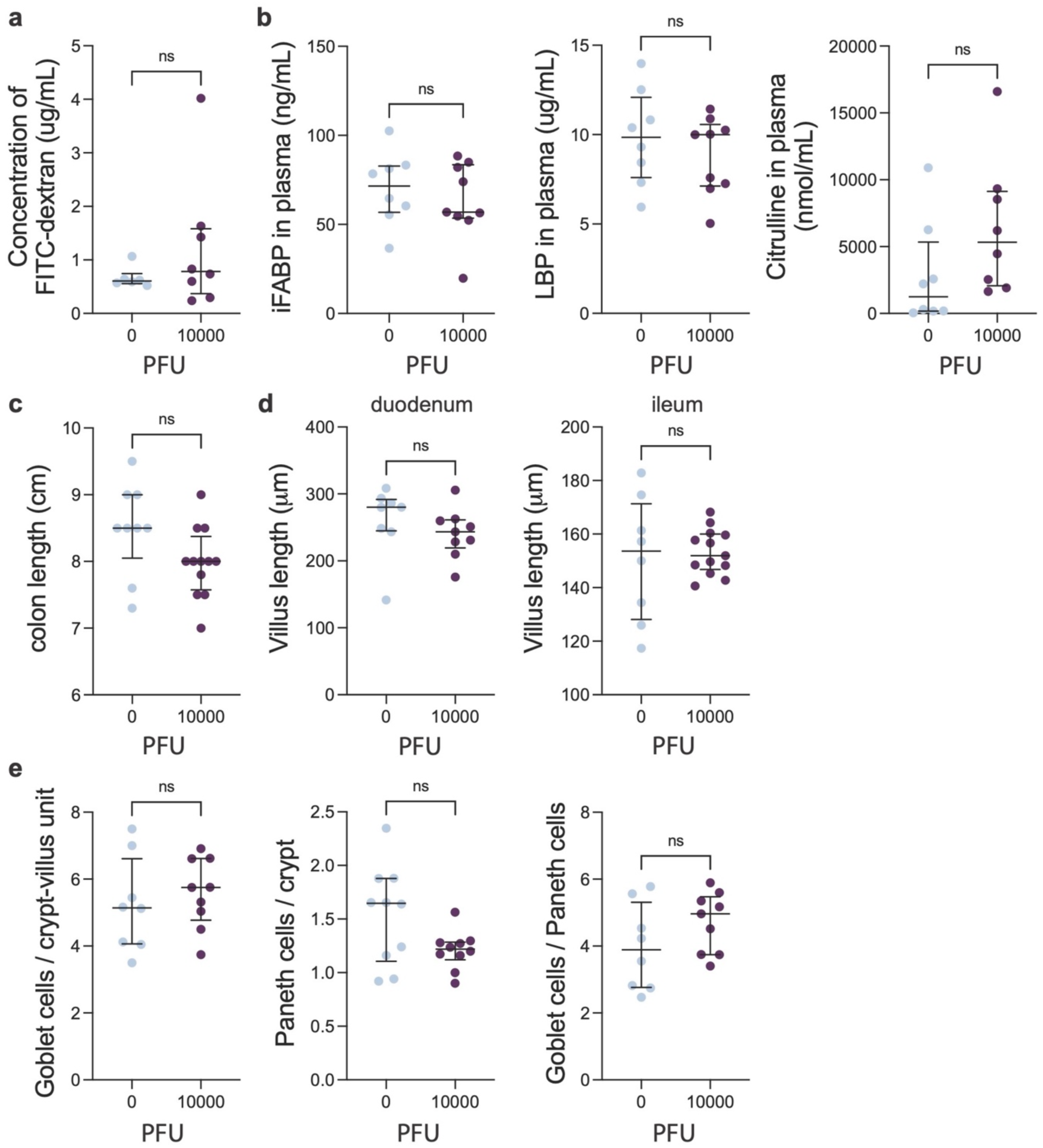
Some intestinal parameters are not modified during SARS-CoV-2 infection. K18-hACE2 mice were analyzed on day 5-6 post intranasal inoculation with 10000 PFU SARS-CoV-2 or mock treatment. **a.** Quantification of fluorescence intensity in the blood following oral administration of FITC-dextran. **B.** Intestinal fatty acid-binding protein (iFABP), LPS-binding protein (LBP), and citrulline concentration in plasma. **C.** Quantification of colon length. **d.** Quantification of villus length in the duodenum (left) and ileum (right) based on H&E staining. **E.** Quantification of goblet cell number (left) and Paneth cell number (middle) per crypt-villus unit in the proximal duodenum based on H&E staining and calculation of goblet cell per Paneth cell ratio based on these quantifications (right). Individual mice, represented by the circles as well as the median and interquartile ranges are depicted. In d, e, each circle shows the mean for each mouse of the cell number counted per crypt-villus unit on 50 units. Results were pooled from 2 (for a) or 3 independent experiments with n=3-5 mice per group for each experiment. Significant differences were determined using the Mann-Whitney U test (ns=non-significant, p > 0.05; **, p < 0.01; ***, p < 0.001; ****, p < 0.0001).

**Extended Data Fig. S4.**
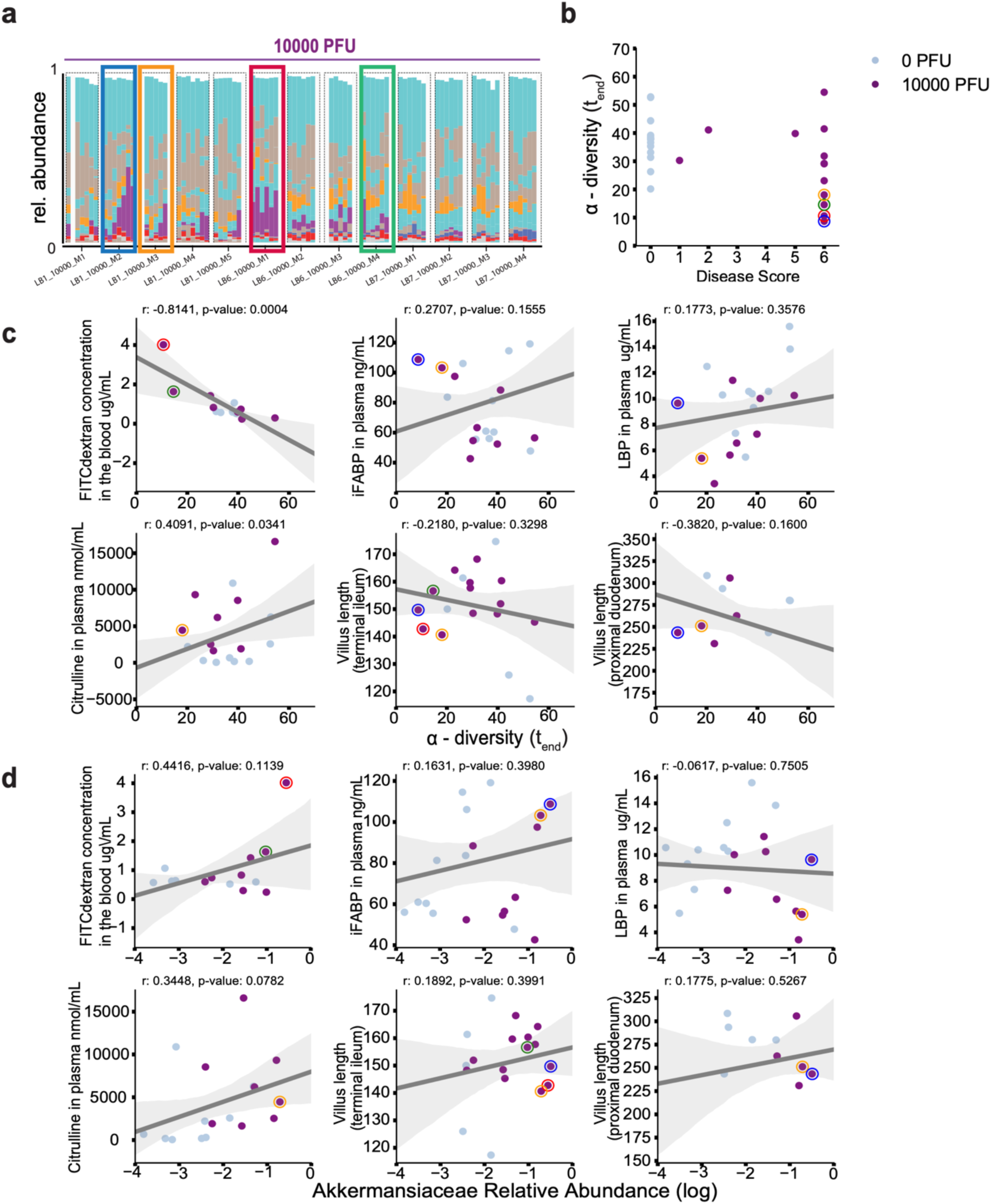
Strongest gut dysbiosis is correlated with markers of defects in the intestinal barrier and epithelium. **a.** Reproduction of Fig. 1 showing bacterial compositions in mice infected with 10^4^ PFUs, highlighting four mice time courses of mice with lowest diversity and highest disease scores at the end of the experiment (**b**). **c-d** Correlations between alpha diversity (**c)** (inverse Simpson) and log_10_ relative *Akkermansia* abundances (**d**) at the end of the experiment with epithelium phenotypes and gut barrier integrity markers measured in the blood of mice (data from mice highlighted in **a** with circles in corresponding colors, lines: linear regression, shaded region: 95%CI).

**Extended Data Fig. S5.**
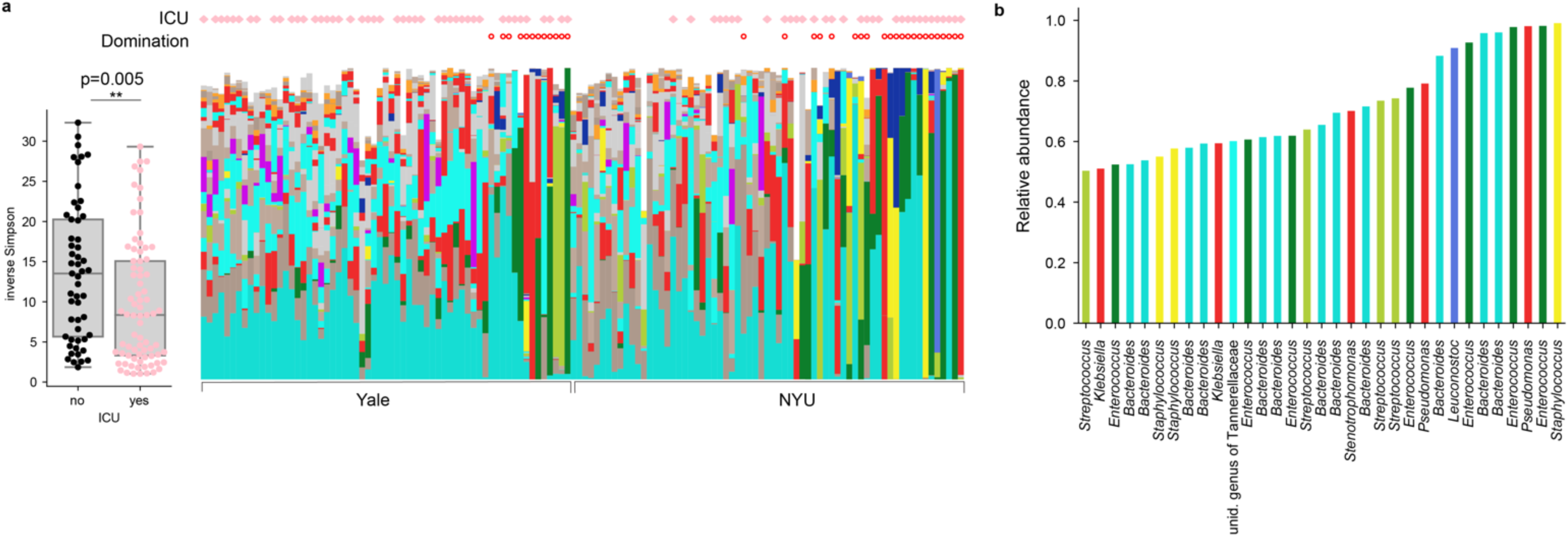
**a** Samples from patients requiring ICU transfer have lower diversity on average (p=0.005, Wilcoxon rank-sum); bars as in Fig. 1 with ICU status of patients and domination state of samples indicated. **b** Genus abundances in samples with a single genus >50% relative abundance.

**Extended Data Fig. S6.**
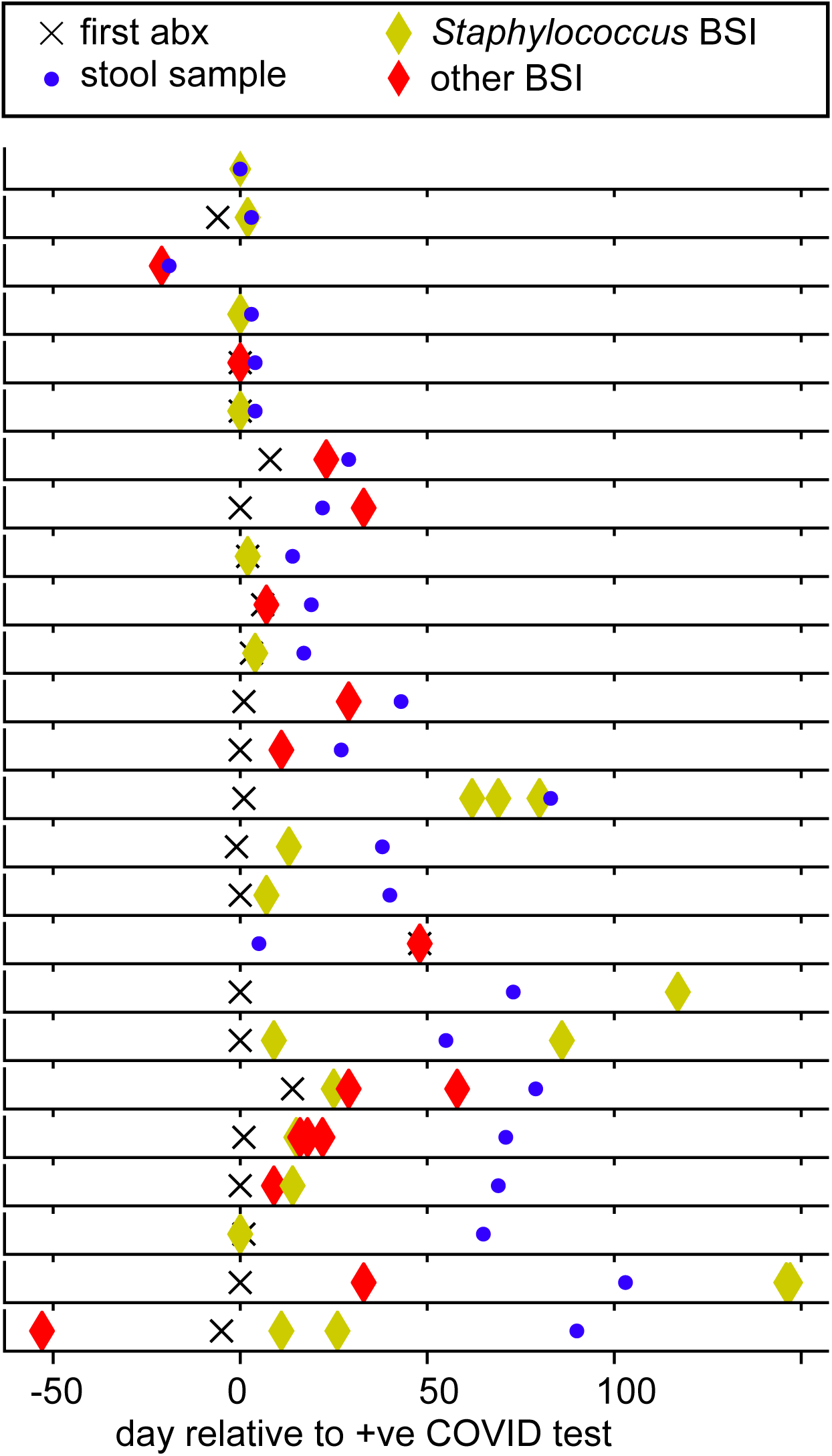
Patients with a positive clinical blood culture result (BSI) received antibiotics, prior or on the day of blood culture results (cross symbol: first recorded antibiotic administration, blue: sequenced stool sample, diamond: positive blood culture result (BSI)).

**Extended Data Fig. S7.**
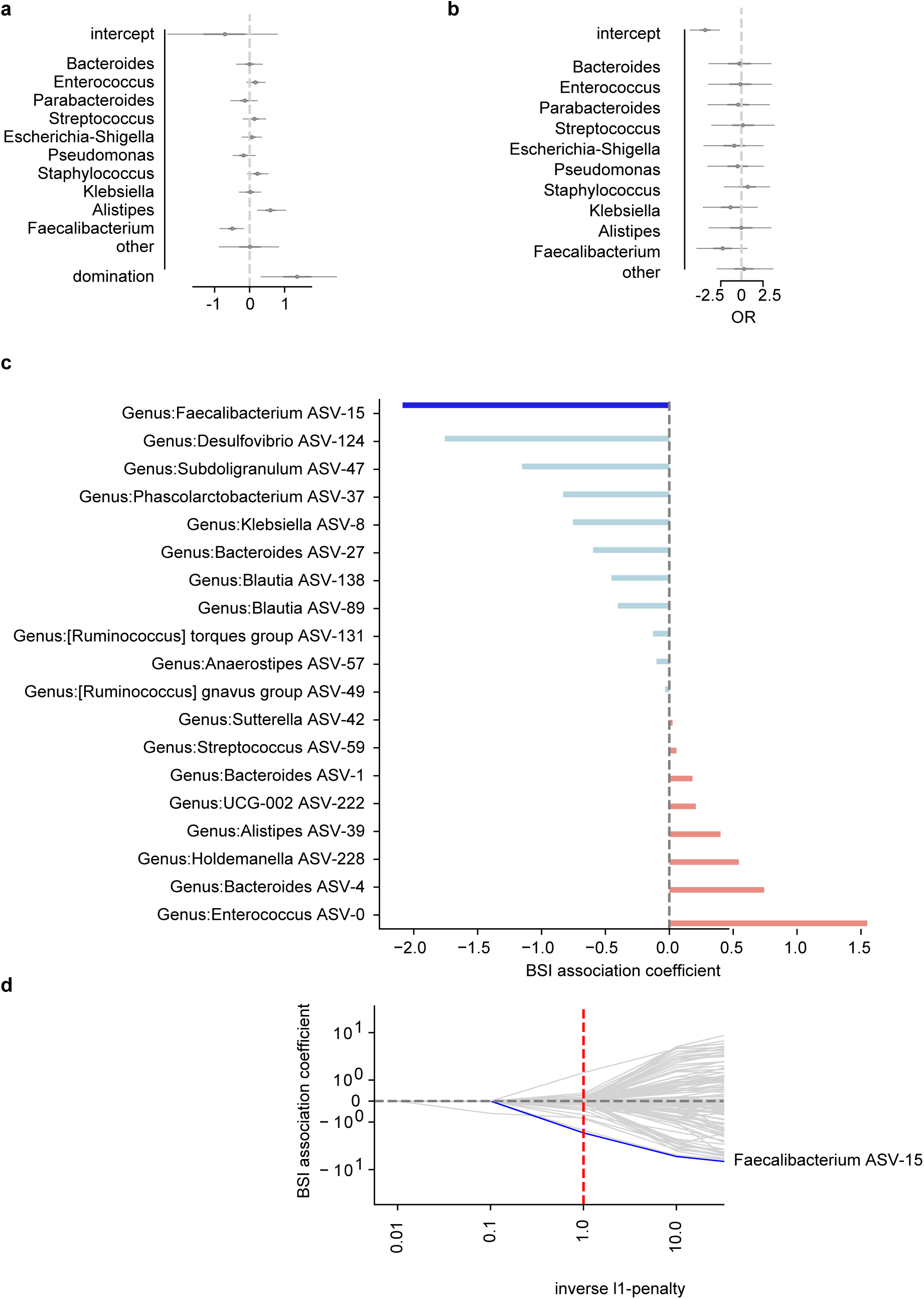
**a** Posterior coefficient estimates from a Bayesian logistic regression regressing log_10_ relative abundances of the top 10 most abundant bacterial genera on BSI status using only BSI cases with associated stool samples taken prior or on the day of a confirmed positive blood culture. **b** Posterior coefficient estimates from a Bayesian logistic regression regressing log_10_ relative abundances of the top 10 most abundant bacterial genera on BSI status with domination status of the microbiome as an additional predictor (domination: >50% of the composition by one taxon). **c** ASVs associated with samples from patients with BSI. Coefficients from a cross-validated, L1-penalized logistic regression correlating the binary outcome (BSI) with log_10_-transformed relative ASV abundances. **d** Cross-validation paths; for all regularization strengths (L1-penalty) used, a *Faecalibacterium* ASV was most negatively associated with BSI-positive samples.

**Extended Data Fig. S8.**
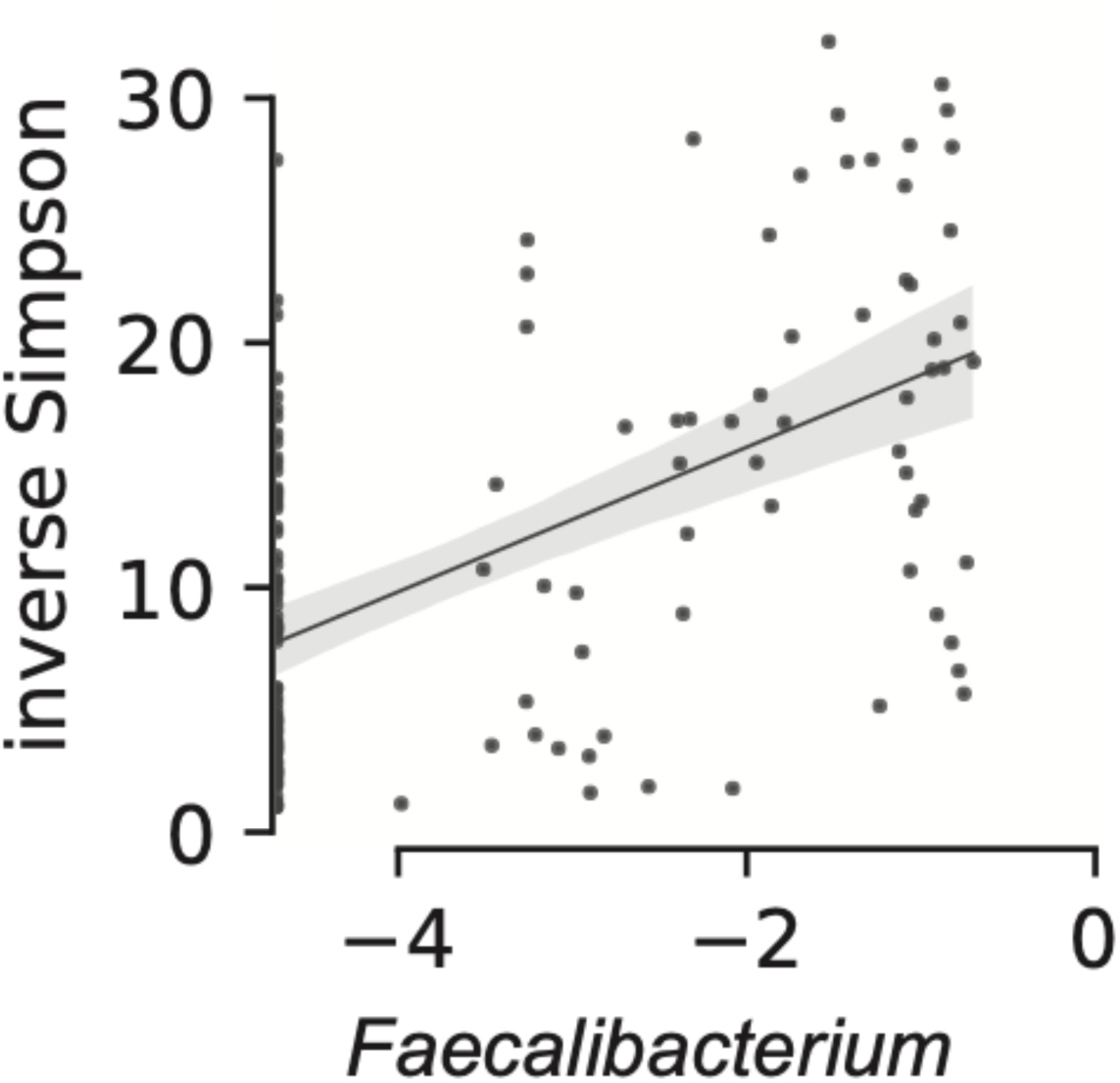
*Faecalibacterium* relative abundance is positively correlated with bacterial alpha diversity. Log10 transformed relative abundances of the genus *Faecalibacterium* in stool samples from patients are correlated with the inverse Simpson diversity index; line from linear regression, shaded region: 95%CI.

**Extended Data Fig. S9.**
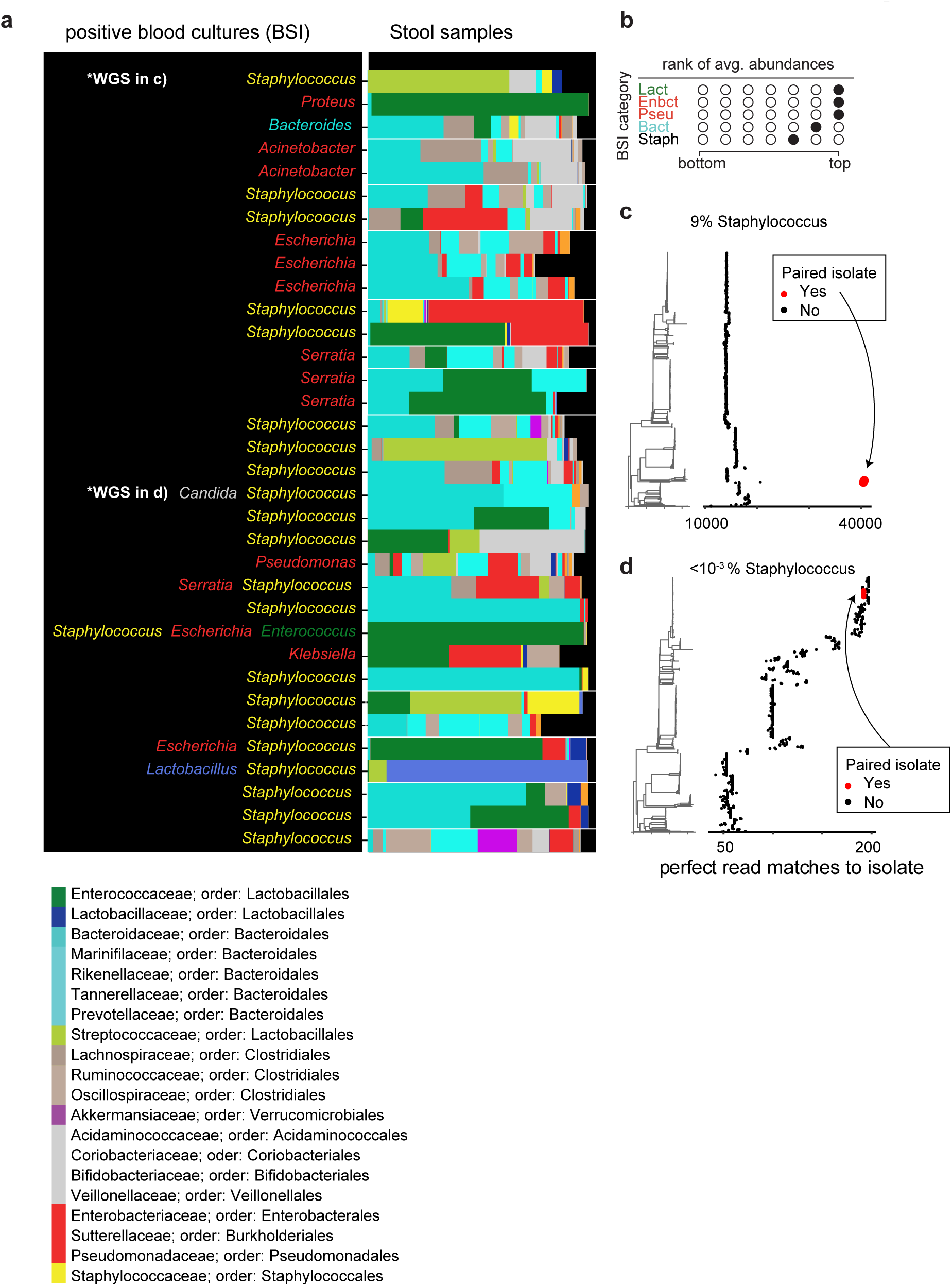
Bacteria in stool of COVID-19 patients match taxa identified blood cultures. **a.** Organisms identified in blood cultures together with bars representing the bacterial family compositions in stool samples; multiple samples belonging to the same patient grouped by a white box. Two samples with matching whole genome sequenced (WGS) blood isolates indicated. **b** Rank analysis of abundance patterns in stool samples from different BSI categories; a filled circle indicates the calculated rank of the focal BSI category (row) in terms of the corresponding taxon stool abundance relative to samples from other BSI categories (Lact: Lactobacillales, Enbct: Enterobacterales; Pseu: Pseudomonadales, Bact: Bacteroidales, Staph: Staphylococcales. Only 5 out of 7 BSI categories are shown because fungal BSIs and the uninfected category have no corresponding bacterial stool abundances). **c,d** left: neighbor-joining tree constructed from all NCBI RefSeq assemblies of *Staphylococcus aureus* genomes in addition to isolates that were isolated from subjects highlighted in **a**. right: counts of perfect read matches of shotgun metagenomic reads from stool samples, red: stool sample sequencing read matches to WGS of isolates from the same patient, black: matches to other genomes.

**Supplementary Table 1:** Clinical characteristics of patients with confirmed COVID-19 at NYU Langone Health and Yale New Haven Hospital.

|  | **NYU, N = 60** | **YALE, N = 36** 926 |
| --- | --- | --- |
| Age (years) | 51 ± 17.5 | 62.52 ± 19.72 |
| Sex (F \| M) | 42% \| 58% | 39% \| 61% |
| **Hospital course and Outcomes** |  |  |
| ICU Admission | 53% | 65% |
| Pneumonia | 42% | 77% |
| Diarrhea | 13% | 32% |
| Intubation | 36% | 41% |
| Sepsis | 23% | 18% |
| Encephalopathy | 12% | 3% |
| Death | 5% | 21% |
| Length of stay (median, IQR) | 37 (10-86) | 27 (11-35.25) |
| **Risk Factors** |  |  |
| Cancer within 1 year | 7% | 4% |
| Chronic Heart Disease | 18% | 36% |
| Hypertension | 38% | 64% |
| Chronic Lung Disease | 7% | 20% |
| Immunosuppression | 17% | 4% |

**Supplementary Table 2:** Clinical characteristics of COVID-19 patients at NYU Langone Health and Yale New Haven Hospital with and without positive blood culture results (BSI).

|  | **BSI, N = 26** | **non-BSI N = 53** |
| --- | --- | --- |
| **Hospital course and Outcomes** |  |  |
| ICU Admission | 69% | 64% |
| Pneumonia | 73% | 53% |
| Diarrhea | 31% | 64% |
| Intubation | 58% | 36% |
| Sepsis | 35% | 21% |
| Encephalopathy | 19% | 6% |
| Death | 15% | 9% |
| Length of stay (median, IQR) | 59 (23-91.5) | 22 (6-51) |

**Supplementary Table 3:** Shotgun metagenomic reads mapped to species identified in clinical blood cultures. Dark grey shading: no sequencing reads from stool samples matched the species identified in clinical blood samples, light grey shading: species of the same genus but not the same species had non-zero read counts in stool samples. The relative abundance of identified species were contrasted with their mean abundances (log10 ratio).

| Organism identified in blood | species identified in stool sample | Log ratio |
| --- | --- | --- |
| *Bacteroides thetaiotaomicron* | Bacteroides_thetaiotaomicron_14-106904-2 | 2.92 |
| *Enterococcus faecalis* Group D | Enterococcus_faecalis_LD33 | 1.8 |
| *Escherichia coli* | Escherichia_coli_K-12_substr._W3110 | 2.2 |
| *Escherichia coli* | Escherichia_coli_IAI39 | 1.6 |
| *Escherichia coli* | Escherichia_coli_536 | 2.8 |
| *Klebsiella pneumoniae* | Klebsiella_pneumoniae_KPNIH27 | -1.7 |
| *Lactobacillus species* | Lactobacillus_curvatus_WiKim38 | 3.9 |
| *Pseudomonas aeruginosa* | Pseudomonas_aeruginosa_SJTD-1 | 3.3 |
| *Serratia marcescens* | Serratia_marcescens_CAV1492 | -0.2 |
| *Staphylococcus aureus* | Staphylococcus_aureus_RF122 | 1.9 |
| *Proteus mirabilis* | Proteus_mirabilis;t_Proteus_mirabilis_BB2000 | 0.56 |
| *Acinetobacter lwolfii* | Acinetobacter_calcoaceticus_EGD_AQ_BF14 | -0.6 |
| *Staphylococcus* | Staphylococcus_sp._HMSC063G01_HMSC063G01 | 0.9 |
| *Staphylococcus* | Staphylococcus_epidermidis_W23144 | 3.3 |
| *Staphylococcus aureus* | not found |  |
| *Staphylococcus hominis* | not found |  |
| *Staphylococcus capitis* | not found |  |
| *Staphylococcus epidermidis, hominis* | Staphylococcus_pseudintermedius_063228 | 2.2 |
| *Staphylococcus epidermidis, hominis ssp hominis* | not found |  |
| *Staphylococcus epidermidis* | Staphylococcus_aureus_JKD6008 | 2.7 |
| *Staphylococcus epidermidis* | Staphylococcus_epidermidis_DAR1907 | 1.1 |
| *Staphylococcus capitis* | Staphylococcus_sp._HMSC067F07_HMSC067F07 | 3.7 |
| *Staphylococcus epidermidis, hominis ssp hominis* | Staphylococcus_hominis_793_SHAE | 0.8 |
| *Staphylococcus epidermidis* | Staphylococcus_sp._HMSC070D05_HMSC070D05 | 3.8 |
| *Staphylococcus hominis, epidermidis* | Staphylococcus_hominis_MMP2 | 1.0 |
| *Staphylococcus hominis, epidermidis* | Staphylococcus_epidermidis_ATCC12228_GCF7645.1 | 0.2 |

**Supplementary Table 4:** SRA accession numbers for the bioproject PRJNA745367 corresponding to the mouse sequencing data. (Excel sheet)

**Supplementary Table 5:** SRA accession numbers for the bioproject PRJNA746322 corresponding to the human stool samples sequencing data. (Excel sheet)

## Bibliography

1. Fajgenbaum, D. C. & June, C. H. Cytokine Storm. N. Engl. J. M ed. 383, 2255–2273 (2020).

2. Lucas, C. et al. Longitudinal analyses reveal immunological misfiring in severe COVID-19. Nature 584, 463–469 (2020).

3. Zuo, T. et al. Alterations in Gut Microbiota of Patients With COVID-19 During Time of Hospitalization. Gastroenterology 159, 944–955.e8 (2020).

4. Yeoh, Y. K. et al. Gut microbiota composition reflects disease severity and dysfunctional immune responses in patients with COVID-19. Gut 70, 698–706 (2021).

5. Gu, S. et al. Alterations of the gut microbiota in patients with coronavirus disease 2019 or H1N1 influenza. Clin. Infect. Dis. 71, 2669–2678 (2020).

6. Nori, P. et al. Bacterial and fungal coinfections in COVID-19 patients hospitalized during the New York City pandemic surge. Infect. Control Hosp. Epidemiol. 42, 84–88 (2021).

7. Grasselli, G. et al. Hospital-Acquired Infections in Critically Ill Patients With COVID-19. Chest (2021). doi:10.1016/j.chest.2021.04.002

8. Yu, D. et al. Low prevalence of bloodstream infection and high blood culture contamination rates in patients with COVID-19. PLoS One 15, e0242533 (2020).

9. Langford, B. J. et al. Bacterial co-infection and secondary infection in patients with COVID-19: a living rapid review and meta-analysis. Clin. Microbiol. Infect. 26, 1622–1629 (2020).

10. Shafran, N. et al. Secondary bacterial infection in COVID-19 patients is a stronger predictor for death compared to influenza patients. Sci. Rep. 11, 12703 (2021).

11. Buffie, C. G. et al. Precision microbiome reconstitution restores bile acid mediated resistance to Clostridium difficile. Nature 517, 205–208 (2015).

12. Buffie, C. G. & Pamer, E. G. Microbiota-mediated colonization resistance against intestinal pathogens. Nat. Rev. Immunol. 13, 790–801 (2013).

13. Modi, S. R., Collins, J. J. & Relman, D. A. Antibiotics and the gut microbiota. J. Clin. Invest. 124, 4212–4218 (2014).

14. Shimasaki, T. et al. Increased Relative Abundance of Klebsiella pneumoniae Carbapenemase-producing Klebsiella pneumoniae Within the Gut Microbiota Is Associated With Risk of Bloodstream Infection in Long-term Acute Care Hospital Patients. Clin. Infect. Dis. 68, 2053–2059 (2019).

15. Kim, S., Covington, A. & Pamer, E. G. The intestinal microbiota: Antibiotics, colonization resistance, and enteric pathogens. Immunol. Rev. 279, 90–105 (2017).

16. Morjaria, S. et al. Antibiotic-Induced Shifts in Fecal Microbiota Density and Composition during Hematopoietic Stem Cell Transplantation. Infect. Immun. 87, (2019).

17. Niehus, R. et al. Quantifying antibiotic impact on within-patient dynamics of extended-spectrum beta-lactamase resistance. Elife 9, (2020).

18. Taur, Y. et al. Intestinal domination and the risk of bacteremia in patients undergoing allogeneic hematopoietic stem cell transplantation. Clin. Infect. Dis. 55, 905–914 (2012).

19. Taur, Y. et al. Reconstitution of the gut microbiota of antibiotic-treated patients by autologous fecal microbiota transplant. Sci. Transl. Med. 10, (2018).

20. Liao, C. et al. Compilation of longitudinal microbiota data and hospitalome from hematopoietic cell transplantation patients. Sci. Data 8, 71 (2021).

21. Peled, J. U. et al. Microbiota as Predictor of Mortality in Allogeneic Hematopoietic-Cell Transplantation. N. Engl. J. Med. 382, 822–834 (2020).

22. McCullers, J. A. The co-pathogenesis of influenza viruses with bacteria in the lung. Nat. Rev. Microbiol. 12, 252–262 (2014).

23. Wang, D. et al. Clinical Characteristics of 138 Hospitalized Patients With 2019 Novel Coronavirus-Infected Pneumonia in Wuhan, China. JAMA 323, 1061–1069 (2020).

24. Westblade, L. F., Simon, M. S. & Satlin, M. J. Bacterial coinfections in coronavirus disease 2019. Trends Microbiol. 29, 930–941 (2021).

25. Sepulveda, J. et al. Bacteremia and Blood Culture Utilization during COVID-19 Surge in New York City. J. Clin. Microbiol. 58, (2020).

26. Lansbury, L., Lim, B., Baskaran, V. & Lim, W. S. Co-infections in people with COVID-19: a systematic review and meta-analysis. J. Infect. 81, 266–275 (2020).

27. Sieswerda, E. et al. Recommendations for antibacterial therapy in adults with COVID-19 - an evidence based guideline. Clin. Microbiol. Infect. 27, 61–66 (2021).

28. Zhai, B. et al. High-resolution mycobiota analysis reveals dynamic intestinal translocation preceding invasive candidiasis. Nat. Med. 26, 59–64 (2020).

29. Haak, B. W. et al. Impact of gut colonization with butyrate-producing microbiota on respiratory viral infection following allo-HCT. Blood 131, 2978–2986 (2018).

30. Deriu, E. et al. Influenza Virus Affects Intestinal Microbiota and Secondary Salmonella Infection in the Gut through Type I Interferons. PLoS Pathog. 12, e1005572 (2016).

31. Yildiz, S., Mazel-Sanchez, B., Kandasamy, M., Manicassamy, B. & Schmolke, M. Influenza A virus infection impacts systemic microbiota dynamics and causes quantitative enteric dysbiosis. Microbiome 6, 9 (2018).

32. Steed, A. L. et al. The microbial metabolite desaminotyrosine protects from influenza through type I interferon. Science 357, 498–502 (2017).

33. Abt, M. C. et al. Commensal bacteria calibrate the activation threshold of innate antiviral immunity. Immunity 37, 158–170 (2012).

34. Ichinohe, T. et al. Microbiota regulates immune defense against respiratory tract influenza A virus infection. Proc. Natl. Acad. Sci. USA 108, 5354–5359 (2011).

35. Sencio, V. et al. Influenza infection impairs the gut’s barrier properties and favors secondary enteric bacterial infection through reduced production of short-chain fatty acids. Infect. Immun. (2021). doi:10.1128/IAI.00734-20

36. Wang, J. et al. Respiratory influenza virus infection induces intestinal immune injury via microbiota-mediated Th17 cell-dependent inflammation. J. Exp. Med. 211, 2397–2410 (2014).

37. Winkler, E. S. et al. SARS-CoV-2 Causes Lung Infection without Severe Disease in Human ACE2 Knock-In Mice. J. Virol. 96, e0151121 (2022).

38. Yinda, C. K. et al. K18-hACE2 mice develop respiratory disease resembling severe COVID-19. PLoS Pathog. 17, e1009195 (2021).

39. Zheng, J. et al. COVID-19 treatments and pathogenesis including anosmia in K18-hACE2 mice. Nature 589, 603–607 (2021).

40. Golden, J. W. et al. Human angiotensin-converting enzyme 2 transgenic mice infected with SARS-CoV-2 develop severe and fatal respiratory disease. JCI Insight 5, (2020).

41. Cadwell, K. et al. A key role for autophagy and the autophagy gene Atg16l1 in mouse and human intestinal Paneth cells. Nature 456, 259–263 (2008).

42. Cadwell, K. et al. Virus-plus-susceptibility gene interaction determines Crohn’s disease gene Atg16L1 phenotypes in intestine. Cell 141, 1135–1145 (2010).

43. Matsuzawa-Ishimoto, Y. et al. Autophagy protein ATG16L1 prevents necroptosis in the intestinal epithelium. J. Exp. Med. 214, 3687–3705 (2017).

44. Schluter, J. et al. The gut microbiota is associated with immune cell dynamics in humans. Nature 588, 303–307 (2020).

45. Gopalakrishnan, V. et al. Gut microbiome modulates response to anti-PD-1 immunotherapy in melanoma patients. Science 359, 97–103 (2018).

46. Diefenbach, C. S. et al. Microbial dysbiosis is associated with aggressive histology and adverse clinical outcome in B-cell non-Hodgkin lymphoma. Blood Adv. 5, 1194–1198 (2021).

47. Sokol, H. et al. Faecalibacterium prausnitzii is an anti-inflammatory commensal bacterium identified by gut microbiota analysis of Crohn disease patients. Proc. Natl. Acad. Sci. USA 105, 16731–16736 (2008).

48. Wrzosek, L. et al. Bacteroides thetaiotaomicron and Faecalibacterium prausnitzii influence the production of mucus glycans and the development of goblet cells in the colonic epithelium of a gnotobiotic model rodent. BMC Biol. 11, 61 (2013).

49. Seibert, B. et al. Mild and Severe SARS-CoV-2 Infection Induces Respiratory and Intestinal Microbiome Changes in the K18-hACE2 Transgenic Mouse Model. Microbiol. Spectr. 9, e0053621 (2021).

50. Sencio, V. et al. Alteration of the gut microbiota following SARS-CoV-2 infection correlates with disease severity in hamsters. Gut Microbes 14, 2018900 (2022).

51. Sokol, H. et al. SARS-CoV-2 infection in nonhuman primates alters the composition and functional activity of the gut microbiota. Gut Microbes 13, 1–19 (2021).

52. Zhang, F. et al. Prolonged Impairment of Short-Chain Fatty Acid and L-Isoleucine Biosynthesis in Gut Microbiome in Patients With COVID-19. Gastroenterology 162, 548– 561.e4 (2022).

53. Gaebler, C. et al. Evolution of antibody immunity to SARS-CoV-2. Nature 591, 639–644 (2021).

54. Park, S.-K. et al. Detection of SARS-CoV-2 in Fecal Samples From Patients With Asymptomatic and Mild COVID-19 in Korea. Clin. Gastroenterol. Hepatol. 19, 1387– 1394.e2 (2021).

55. Xiao, F. et al. Evidence for Gastrointestinal Infection of SARS-CoV-2. Gastroenterology 158, 1831–1833.e3 (2020).

56. Cheung, K. S. et al. Gastrointestinal Manifestations of SARS-CoV-2 Infection and Virus Load in Fecal Samples From a Hong Kong Cohort: Systematic Review and Meta-analysis. Gastroenterology 159, 81–95 (2020).

57. Lamers, M. M. et al. SARS-CoV-2 productively infects human gut enterocytes. Science 369, 50–54 (2020).

58. Cao, J. et al. Integrated gut virome and bacteriome dynamics in COVID-19 patients. Gut Microbes 13, 1–21 (2021).

59. Klag, T., Stange, E. F. & Wehkamp, J. Defective antibacterial barrier in inflammatory bowel disease. Dig. Dis. 31, 310–316 (2013).

60. Ramanan, D. & Cadwell, K. Intrinsic defense mechanisms of the intestinal epithelium. Cell Host Microbe 19, 434–441 (2016).

61. Schluter, J. & Foster, K. R. The evolution of mutualism in gut microbiota via host epithelial selection. PLoS Biol. 10, e1001424 (2012).

62. McLoughlin, K., Schluter, J., Rakoff-Nahoum, S., Smith, A. L. & Foster, K. R. Host selection of microbiota via differential adhesion. Cell Host Microbe 19, 550–559 (2016).

63. Fernandez-Castañer, M. et al. Evaluation of B-cell function in diabetics by C-peptide determination in basal and postprandial urine. Diabete Metab 13, 538–542 (1987).

64. Yu, S. et al. Paneth Cell-Derived Lysozyme Defines the Composition of Mucolytic Microbiota and the Inflammatory Tone of the Intestine. Immunity 53, 398–416.e8 (2020).

65. Salzman, N. H. et al. Enteric defensins are essential regulators of intestinal microbial ecology. Nat. Immunol. 11, 76–83 (2010).

66. van der Lugt, B. et al. Akkermansia muciniphila ameliorates the age-related decline in colonic mucus thickness and attenuates immune activation in accelerated aging Ercc1-/Δ7 mice. Immun. Ageing 16, 6 (2019).

67. Wang, L. et al. An observational cohort study of bacterial co-infection and implications for empirical antibiotic therapy in patients presenting with COVID-19 to hospitals in North West London. J. Antimicrob. Chemother. 76, 796–803 (2021).

68. Labarta-Bajo, L. et al. Type I IFNs and CD8 T cells increase intestinal barrier permeability after chronic viral infection. J. Exp. Med. 217, (2020).

69. Karki, R. et al. Synergism of TNF-α and IFN-γ Triggers Inflammatory Cell Death, Tissue Damage, and Mortality in SARS-CoV-2 Infection and Cytokine Shock Syndromes. Cell 184, 149–168.e17 (2021).

70. Giron, L. B. et al. Plasma Markers of Disrupted Gut Permeability in Severe COVID-19 Patients. Front. Immunol. 12, 686240 (2021).

71. Xie, X. et al. An Infectious cDNA Clone of SARS-CoV-2. Cell Host Microbe 27, 841– 848.e3 (2020).

72. Gohl, D. M. et al. Systematic improvement of amplicon marker gene methods for increased accuracy in microbiome studies. Nat. Biotechnol. 34, 942–949 (2016).

73. Wang, Q., Garrity, G. M., Tiedje, J. M. & Cole, J. R. Naive Bayesian classifier for rapid assignment of rRNA sequences into the new bacterial taxonomy. Appl. Environ. Microbiol. 73, 5261–5267 (2007).

74. Pawlowsky-Glahn, V., Egozcue, J. J. & Tolosana-Delgado, R. *Modelling and analysis of compositional data*. (John Wiley & Sons, Ltd, 2015). doi:10.1002/9781119003144

75. Kruschke, J. K. Bayesian estimation supersedes the t test. J. Exp. Psychol. Gen. 142, 573– 603 (2013).

76. Homan, M. D. & Gelman, A. The No-U-Turn Sampler: Adaptively Setting Path Lengths in Hamiltonian Monte Carlo. J. Mach. Learn. Res. 15, 1593–1623 (2014).

